# Nitric oxide is not responsible for the initial sensory-induced neurovascular coupling response in mouse cortex

**DOI:** 10.1101/2022.05.24.493260

**Authors:** L Lee, L Boorman, E Glendenning, C Shen, J Berwick, C Howarth

## Abstract

Neurovascular coupling ensures that changes in neural activity are accompanied by localised changes in cerebral blood flow. While much is known about the involvement of excitatory neurons in neurovascular coupling, the role of inhibitory interneurons is unresolved. While nNOS-expressing interneurons have been shown to be capable of eliciting vasodilation, the role of nitric oxide in functional hyperemia remains a matter of debate. Therefore in the present study we applied a combination of optogenetic and pharmacological approaches, 2-dimensional optical imaging spectroscopy, and electrophysiology to investigate the role of nitric oxide in neurovascular coupling responses evoked by nNOS-expressing interneurons and whisker stimulation in mouse sensory cortex. The haemodynamic response evoked by nNOS-expressing interneurons was significantly altered in the presence of the NOS inhibitor LNAME, revealing a large initial 20-HETE-dependent vasoconstriction. In contrast, the haemodynamic response induced by sensory stimulation was largely unchanged by LNAME. Our results suggest that while nitric oxide plays a key role in neurovascular responses evoked by nNOS-expressing interneurons it does not mediate the initial sensory-induced neurovascular coupling response in mouse cortex. Thus, our results call into question the involvement of nNOS-expressing interneurons and nitric oxide in sensory-evoked functional hyperemia.

## Introduction

The ability to regulate cerebral blood flow (CBF) in a localised and dynamic manner in response to neuronal activity is essential for the maintenance of healthy brain function. While the regulation of CBF by glutamatergic neurons has been well investigated (Attwell et al., 2010), regulation by GABAergic inhibitory interneurons (INs) has received less attention. Using optogenetic approaches to specifically activate distinct neural populations, recent research has revealed not only that cortical INs are capable of regulating CBF (Anenberg et al., 2015; Uhlirova et al., 2016; Vazquez et al., 2018), but that subpopulations of cortical INs, which release different vasoactive molecules (Cauli et al., 2004), can drive distinct haemodynamic responses (Dahlqvist et al., 2020; Lee et al., 2020; Krawchuk et al., 2020; Lee et al., 2021). The vasodilatory pathways involved in specific IN-evoked haemodynamic responses remain understudied. Understanding how, and when, INs regulate CBF would inform our interpretation of functional imaging signals such as blood oxygen level dependent functional magnetic resonance imaging (BOLD fMRI), in which blood-based signals act as a proxy for neural activity (Howarth et al., 2021). Furthermore, understanding IN regulation of CBF has relevance for conditions in which altered CBF and IN dysfunction have previously been reported, such as Alzheimer’s disease (Iturria-Medina et al., 2016; Palop and Mucke, 2016; Verret et al., 2012) , epilepsy (Dudek, 2020; Harris et al., 2013) and ageing (Beishon et al., 2021; Miettinen et al., 1993).

Having previously demonstrated that neuronal nitric oxide synthase-expressing interneurons (nNOS INs) are able to evoke robust changes in cerebral blood volume and oxygenation (Lee et al., 2020), in this study we aimed to investigate the vasodilatory pathways underlying nNOS IN-evoked haemodynamic changes. A prime candidate was nitric oxide (NO), which is produced by three isoforms of the enzyme nitric oxide synthase (NOS): neuronal NOS (nNOS), expressed in neurons; endothelial NOS (eNOS), expressed in vascular endothelial cells; and inducible NOS (iNOS), expressed during inflammatory responses (Förstermann and Sessa, 2012). NO release by nNOS-expressing INs (which target blood vessels (Shaw et al., 2021; Vlasenko et al., 2007)) may underpin CBF responses evoked by VGAT-expressing INs (Vazquez et al., 2018). Furthermore, NO is involved in maintaining basal vascular tone (Echagarruga et al., 2020) and has been suggested as a key mediator in sensory-evoked neurovascular coupling (NVC; recently reviewed by Hosford and Gourine (2019)). NO can evoke vasodilation via several vasoactive pathways. In addition to relaxing smooth muscle cells via a cGMP-associated pathway (Faraci and Brian, 1994), NO can also inhibit the production of 20-hydroxyeicosatetraenoic acid (20-HETE; Oyekan et al., 1999), a vasoconstrictor (Gebremedhin et al., 2000; Harder et al., 2011; Imig et al., 1996), thus enhancing dilation (Alonso-Galicia et al., 1999; Liu et al., 2008; Sun et al., 1998). However, the role of NO in neurovascular coupling remains unresolved, with genetic or pharmacological inhibition of NOS reported to decrease (Ances et al., 2010; Hoiland et al., 2020; Liu et al., 2008; Stefanovic et al., 2007; Toth et al., 2015), increase (Ances et al., 2010; Hariharan et al., 2019) and have no effect on (Ances et al., 2010; Ayata et al., 1996; Liu et al., 2008; Ma et al., 1996; Vazquez et al., 2018; Wang et al., 1993; White et al., 1999) haemodynamic responses to sensory stimulation. Therefore, this study investigated the involvement of NO in nNOS IN- evoked and sensory stimulation-evoked haemodynamic changes. To this end, in whisker barrel cortex of lightly anaesthetised mice, we recorded haemodynamic responses to short duration (2s) optogenetic activation of nNOS INs and whisker stimulation, in the presence and absence of the pharmacological non-selective NOS inhibitor N-nitro-L-arginine-methyl-ester (LNAME), which suppresses NO production. We report that although NO is involved in the initial haemodynamic response to nNOS IN activation, counteracting the constrictive effects of 20-HETE release, NO is not involved in the initiation of sensory-evoked haemodynamic responses but does alter return to baseline dynamics.

## Results

### NO is involved in the initial nNOS IN-evoked haemodynamic response, but not in the initiation of a sensory-evoked haemodynamic response

Genetically modified mice expressing channelrhodopsin-2 (ChR2) in nNOS INs were used to investigate the involvement of nitric oxide (NO) in nNOS IN- and sensory-evoked cortical haemodynamic responses. As previously described, ChR2 expression occurs in the appropriate neurons in these mice with approximately 90% expression (Lee et al., 2020), enabling the use of blue light to selectively activate nNOS INs. In lightly anaesthetised mice we combined 2-dimensional optical imaging spectroscopy (2D-OIS), optogenetics and pharmacological blockade to assess whether the localised haemodynamic response evoked by short duration (2s) nNOS IN activation and sensory stimulation were dependent on NO produced by NOS. High resolution 2D maps of stimulation-evoked changes in blood volume (Hbt), oxygenated haemoglobin (Hbo) and deoxygenated haemoglobin (Hbr) were recorded before and after treatment with LNAME (75mg/kg, i.p. (Bannerman et al., 1994)). Mice first received a short duration (2s) mechanical stimulation of the whiskers to define the whisker barrel cortex and guide placement of the optrode and, where appropriate, multi-channel electrode. The optrode, consisting of a fibre optic-coupled blue LED (470nm) placed directly above the centre of the whisker barrel cortex, delivered the photostimulation necessary to activate ChR2. Each animal received interleaved whisker stimulations (2s, 5Hz), photostimulations (2s, 99Hz) and, in a subset of animals, simultaneous photostimulation and whisker stimulation. The resulting haemodynamic changes were centred around the optrode (photostimulation: Fig 1A) and localised to the whisker barrel cortex (whisker stimulation: Fig 1D), as we have previously reported (Lee et al., 2020). Prior to treatment with LNAME, both 2s photostimulation of cortical nNOS INs (Fig. 1 A,C, top rows) and sensory stimulation (Fig. 1 D,E, top rows) elicited localised increases in concentration of Hbt and Hbo and decreases in Hbr concentration. As previously described (Lee et al., 2020), the largest increases in Hbt and Hbo were observed in branches of the middle cerebral artery overlying the whisker barrel cortex, and a decrease in Hbr was observed in the draining veins (Fig 1A, D, top row). In response to photostimulation of nNOS INs, the time series of the haemodynamic response taken from an arterial region of interest (ROI, Fig. 1B) revealed a bidirectional response comprising of an initial decrease in Hbt and Hbo accompanied by a concomitant increase in Hbr (‘initial dip’), followed by an increase in Hbt and Hbo and decrease in Hbr (‘peak’), which peaked following the cessation of stimulation (Fig. 1F, top left). In contrast, the time series of the haemodynamic response to whisker stimulation consisted solely of an increase in Hbt and Hbo and corresponding washout of Hbr, which peaked following the cessation of stimulation (‘peak’, Fig 1F, bottom left), with no measurable ‘initial dip’.

**Figure 1:**
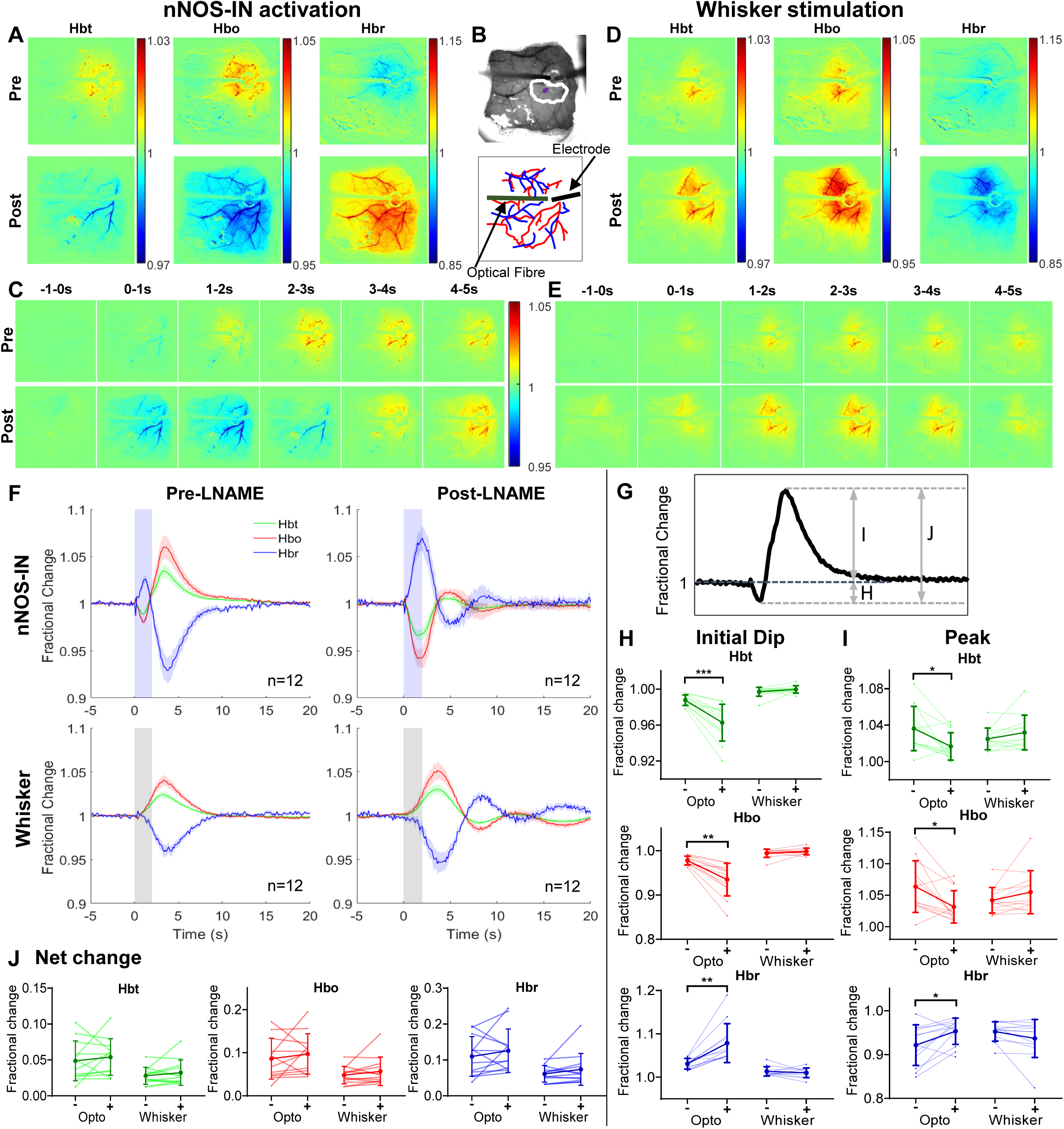
NOS-dependence of haemodynamic responses evoked by 2s nNOS-IN activation or whisker stimulation. **(A-E):** Data from representative mouse showing haemodynamic response to 2s optogenetic stimulation of nNOS-INs **(A-C)** or 2s whisker stimulation **(D-E)**. **(A,D)**: Trial-averaged stimulation-evoked changes in Hbt, Hbo and Hbr, as compared to baseline, pre-LNAME (upper) and post-LNAME (lower) injection. Color bars represent fractional change. **(B):** Thinned cranial window (upper) with cortical surface vasculature visible (imaged at 575nm illumination), optic fibre and electrode can be seen. White ROI indicates whisker barrel cortex and purple ROI indicates arteriole region, from which timeseries data are extracted. Diagram (lower) indicating surface arteries (red) and veins (blue) visible through thinned skull window. **(C,E):** Evolution of trial-averaged changes in Hbt. Stimulation of nNOS-INs **(C)** or whiskers **(E)** occurs at 0-2s. Colour bar represents fractional change. **(F):** Group data (n=12 mice). Mean fractional change in Hbt, Hbo and Hbr in arteriolar ROI before (left) and after (right) LNAME injection. Blue shading indicates photostimulation period (upper), grey shading indicates whisker stimulation period (lower). Data: mean ± SEM **(G):** Hbt time series from example mouse illustrating metrics for analysis: initial dip (H), peak (I), net change (minima to maxima, J). **(H-J):** Fractional change in Hbt, Hbo and Hbr in response to optogenetic and whisker stimulation with (+) and without (-) NOS inhibition by LNAME. Darker lines represent group mean ± SD, lighter lines indicate trial-averaged mean for individual animals. n = 12 mice. *p<0.05, **p<0.01, ***p<0.001. **(H):** Initial fractional change (‘initial dip’). **(I):** Maximum fractional change (‘peak’). **(J):** Maximal stimulation-evoked net change (minima to maxima).

In agreement with reports that cortical NOS activity is reduced by more than 90% one hour after i.p. injection of LNAME (Bannerman et al., 1994), 70 minutes was found to be sufficient time for a significant effect of LNAME to be observed (Fig 2; F(1.361,10.889)=11.65, p=0.004, η^2^=0.593; see Table 1 for pairwise comparisons). Therefore, post-LNAME haemodynamic measurements were obtained 70-135 minutes after systemic injection of LNAME. Following treatment with LNAME, the nNOS IN-evoked haemodynamic response was inverted, showing a decrease in Hbt and Hbo and an increase in Hbr (Fig 1A). Inspection of the timeseries of the haemodynamic response (Fig 1F) revealed a greater reduction in Hbt during the photostimulation period (as compared to pre-LNAME), followed by an increase in Hbt which peaked after stimulation offset (Fig 1C,F). In contrast, the haemodynamic response evoked by whisker stimulation was unchanged by LNAME in terms of polarity and timing (Fig 1D). The localised haemodynamic response to whisker stimulation consisted of an increase in Hbt during the stimulation period, which peaked after stimulation offset, both before and after treatment with LNAME (Fig 1E, F).

**Figure 2:**
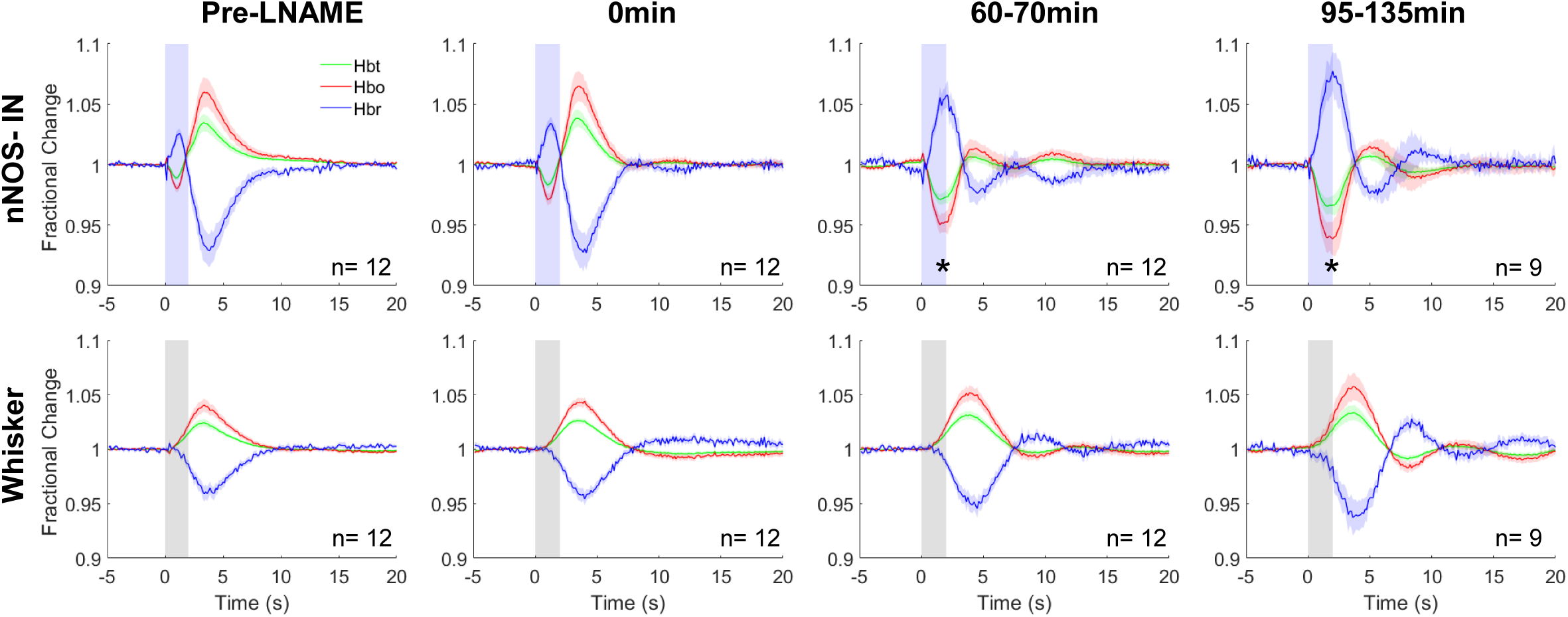
Development of effect of LNAME: Mean fractional change in Hbt, Hbo and Hbr in arteriolar ROI in response to 2s optogenetic stimulation of nNOS-INs (top row) or 2s whisker stimulation (lower row) at different time points relative to injection of LNAME. Column 1: experiment immediately prior to LNAME injection; Column 2: experiment immediately following LNAME injection; Column 3: experiment commencing 60-70 min post LNAME injection; Column 4: experiment commencing 95-135 min post LNAME injection. Blue shading indicates photostimulation period (top row), grey shading indicates whisker stimulation period (lower row). Data are mean ± SEM, n indicates number of mice. * p<0.05 Hbt minima compared to Pre-LNAME.

**Table 1:** Pairwise comparisons of Hbt initial dip evoked by nNOS-IN activation at different time points relative to LNAME injection (Bonferroni adjusted p values reported). (n= 9 mice) * p<0.05

| Pairwise comparison | p |
| --- | --- |
| Pre-LNAME vs 0mins | 0.32 |
| Pre-LNAME vs 60-70mins | 0.02* |
| Pre-LNAME vs 95-135mins | 0.017* |
| 0mins vs 60-70mins | 0.099 |
| 0mins vs 95-135mins | 0.113 |
| 60-70mins vs 95-135mins | 0.798 |

A subset of animals underwent acute implantation of an electrode, to allow concurrent measurement of evoked haemodynamic (Fig 1) and neural (Fig 3) changes. As electrode placement elicits a cortical spreading depolarisation (CSD) which can have long-lasting confounding effects on haemodynamic measures (Shabir et al., 2022; Sharp et al., 2020), insertion of an electrode was included as a factor in the statistical analysis of the haemodynamic data. Therefore, a 3-way mixed ANOVA was used to assess the effect of LNAME, stimulation type, and electrode insertion on the evoked haemodynamic response. Due to the bidirectional nature of the nNOS IN-evoked haemodynamic response, three metrics were analysed (Fig 1G), (i) maximal change during initial response (‘initial dip’, 0.25-5s after stimulation onset, Fig 1H), (ii) maximal change during later response (‘peak’, 0.25-10s after stimulation onset, Fig 1I), and (iii) net change (‘peak’-‘initial dip’, Fig 1J). For all haemodynamic profiles (Hbt, Hbo and Hbr), for all metrics considered, no significant effect of electrode insertion was found (Table 2-4), therefore haemodynamic data from all mice were combined (Fig 1). This lack of effect of CSD on haemodynamic responses is likely due to the fact that electrode insertion occurred approximately 50 minutes prior to collection of pre-LNAME data, sufficient time for haemodynamic recovery after CSD (Chang et al., 2010; Shabir et al., 2022). As a statistically significant interaction between stimulation type and LNAME was revealed for all haemodynamic profiles (initial dip: Table 2, peak: Table 3), suggesting that the effect of LNAME depends on the type of stimulation applied, simple effects tests to assess the effect of LNAME for each stimulation type were performed.

**Figure 3:**
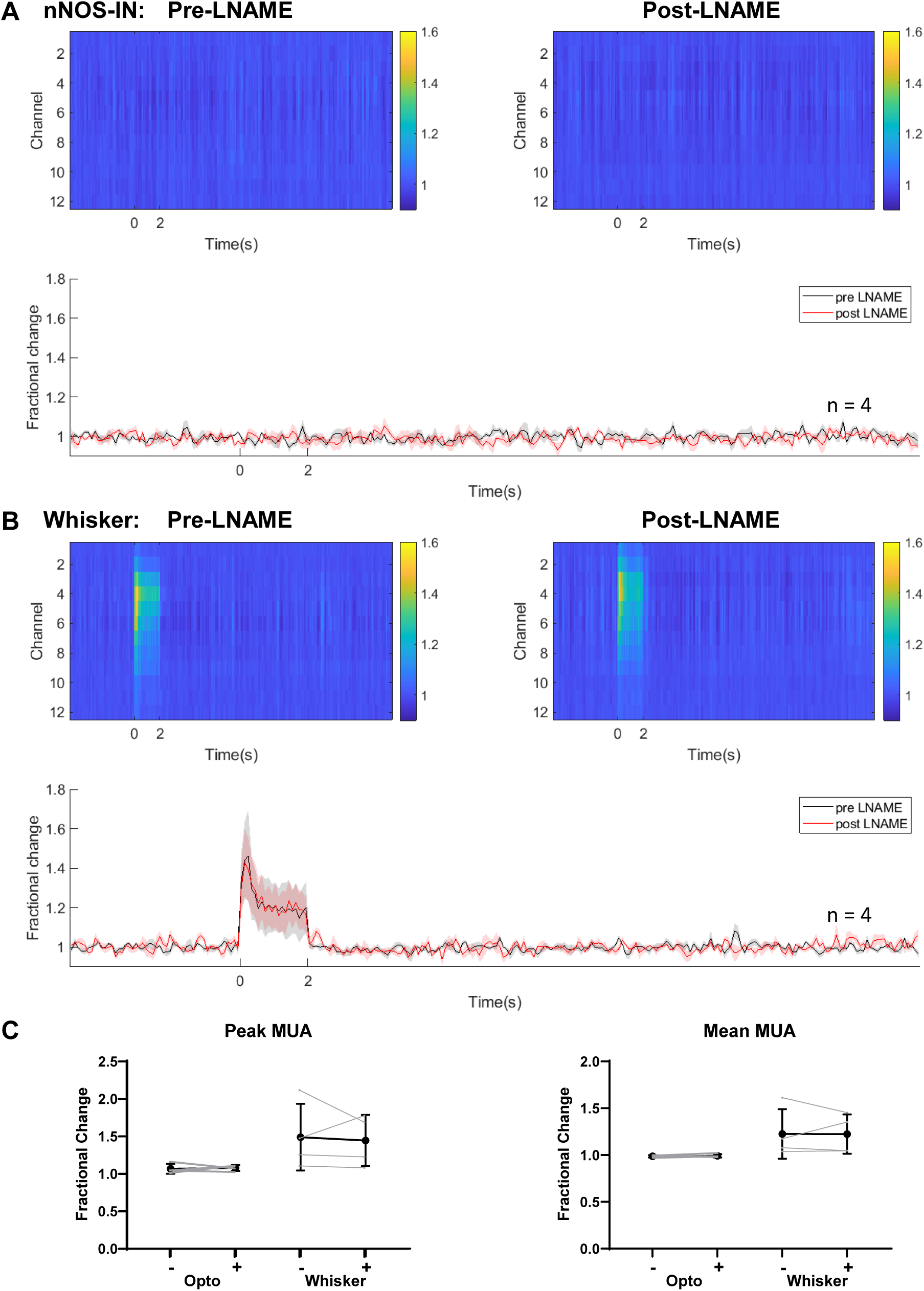
Stimulation-evoked multi-unit activity (MUA) is unchanged in presence of LNAME. Neural responses to **(A):** 2s optogenetic stimulation of nNOS-INs or **(B):** 2s whisker stimulation (stimulations start at 0s). **Top row:** mean change in MUA compared to baseline throughout cortical depth (as indicated by electrode channel number). Colour bar represents fractional change. **Left:** Pre-LNAME injection. **Right:** Post-LNAME injection. **Bottom row:** mean time series of response taken from channels 3-6 of electrode (mean ± SEM). Data were collected concurrently with subset of haemodynamic responses displayed in Figure 1. **(C):** Peak and Mean MUA during 2s optogenetic or whisker stimulation with (+) and without (-) LNAME. Darker lines represent group mean ± SD, lighter lines indicate trial-averaged means for individual animals. n=4 mice for all panels.

**Table 2:** Effect of LNAME on haemodynamic response – Initial Dip: 3-way mixed ANOVA results. Bonferroni correction for multiple comparisons was applied, to account for three different haemodynamic profiles. (n= 12 mice) *p<0.017, **p<0.003, *** p<0.0003.

| Variable | Factor | F(1,10) |  |  |
| --- | --- | --- | --- | --- |
| | | F | p | $\eta^2$ |
| Hbt response in initial dip period | Electrode | 1.208 | 0.297 | 0.108 |
|  | Stim | 31.075 | 0.000*** | 0.757 |
|  | Stim*Electrode | 0.572 | 0.467 | 0.054 |
|  | Drug | 21.2 | 0.001** | 0.679 |
|  | Drug*Electrode | 0.001 | 0.982 | 0.000 |
|  | Stim*Drug | 25.563 | 0.000** | 0.719 |
|  | Stim*Drug*Electrode | 0.137 | 0.719 | 0.014 |
| Hbo response in initial dip period | Electrode | 0.897 | 0.366 | 0.082 |
|  | Stim | 31.342 | 0.000*** | 0.758 |
|  | Stim*Electrode | 0.383 | 0.55 | 0.037 |
|  | Drug | 17.618 | 0.002** | 0.638 |
|  | Drug*Electrode | 0.000 | 0.994 | 0.000 |
|  | Stim*Drug | 22.81 | 0.001** | 0.695 |
|  | Stim*Drug*Electrode | 0.024 | 0.881 | 0.002 |
| Hbr response in initial dip period | Electrode | 0.282 | 0.607 | 0.027 |
|  | Stim | 27.526 | 0.000** | 0.734 |
|  | Stim*Electrode | 0.014 | 0.908 | 0.001 |
|  | Drug | 13.484 | 0.004* | 0.574 |
|  | Drug*Electrode | 0.018 | 0.896 | 0.002 |
|  | Stim*Drug | 16.223 | 0.002** | 0.619 |
|  | Stim*Drug*Electrode | 0.036 | 0.853 | 0.004 |

| Simple effects tests: LNAME (2s whisker stimulation) | p |
| --- | --- |
| Hbt response in initial dip period | 0.159 |
| Hbo response in initial dip period | 0.201 |
| Hbr response in initial dip period | 0.434 |

**Table 3:** Effect of LNAME on haemodynamic response – Peak: 3-way mixed ANOVA results. Bonferroni correction for multiple comparisons was applied, to account for three different haemodynamic profiles. (n= 12 mice) *p<0.017, **p<0.003, *** p<0.0003.

| Variable | Factor | F(1,10) |  |  |
| --- | --- | --- | --- | --- |
| | | F | p | $\eta^2$ |
| Peak Hbt response | Electrode | 2.058 | 0.182 | 0.171 |
|  | Stim | 0.085 | 0.776 | 0.008 |
|  | Stim*Electrode | 0.01 | 0.924 | 0.001 |
|  | Drug | 3.058 | 0.111 | 0.234 |
|  | Drug*Electrode | 2.851 | 0.122 | 0.222 |
|  | Stim*Drug | 9.229 | 0.013* | 0.48 |
|  | Stim*Drug*Electrode | 1.45 | 0.256 | 0.127 |
| Peak Hbo response | Electrode | 1.966 | 0.191 | 0.164 |
|  | Stim | 0.008 | 0.931 | 0.001 |
|  | Stim*Electrode | 0.015 | 0.905 | 0.001 |
|  | Drug | 2.287 | 0.161 | 0.186 |
|  | Drug*Electrode | 2.533 | 0.143 | 0.202 |
|  | Stim*Drug | 8.795 | 0.014* | 0.468 |
|  | Stim*Drug*Electrode | 1.433 | 0.259 | 0.125 |
| Peak Hbr response | Electrode | 2.079 | 0.180 | 0.172 |
|  | Stim | 0.312 | 0.589 | 0.03 |
|  | Stim*Electrode | 0.09 | 0.77 | 0.009 |
|  | Drug | 0.981 | 0.345 | 0.089 |
|  | Drug*Electrode | 1.641 | 0.229 | 0.141 |
|  | Stim*Drug | 8.551 | 0.015* | 0.461 |
|  | Stim*Drug*Electrode | 1.744 | 0.216 | 0.149 |

| Simple effects tests: LNAME (2s whisker stimulation) | p |
| --- | --- |
| Peak Hbt response | 0.129 |
| Peak Hbo response | 0.124 |
| Peak Hbr response | 0.127 |

In the presence of LNAME, nNOS IN stimulation evoked a larger ‘initial dip’ (Fig 1H), resulting in a larger initial decrease in Hbt (pre= 0.988±0.002, post= 0.963±0.006, p=0.000) and Hbo (pre= 0.977±0.003, post= 0.935±0.011, p=0.001), and a greater initial increase in Hbr (pre= 1.031±0.004, post= 1.079±0.014, p= 0.002). Furthermore, post LNAME the ‘peak’ haemodynamic response to nNOS IN activation was reduced across all haemoglobin components (Hbt: pre= 1.036±0.007, post= 1.017±0.005, p= 0.013; Hbo: pre= 1.064±0.011, post= 1.032±0.008, p= 0.016; Hbr: pre= 0.921±0.012, post= 0.953±0.009, p= 0.023, Fig 1I), as compared to before LNAME injection.

The maximum nNOS IN-evoked net change (i.e. minima to maxima) in Hbt, Hbo and Hbr was unchanged in the presence of LNAME (Fig 1J, Table 4). These data suggest that while the initial haemodynamic response to nNOS IN activity is dependent on NO production by NOS, a second, NO-independent, pathway underlies the later increases in Hbt, Hbo and associated washout of Hbr.

**Table 4:** Effect of LNAME on haemodynamic response - net change: 3-way mixed ANOVA results. Bonferroni correction for multiple comparisons was applied, to account for three different haemodynamic profiles. (n=12 mice) *p<0.017, **p<0.003, ***p<0.0003.

| Variable | Factor | F(1,10) |  |  |
| --- | --- | --- | --- | --- |
| | | F | p | $\eta^2$ |
| Maximum net change in Hbt | Electrode | 2.118 | 0.176 | 0.175 |
|  | Stim | 12.499 | 0.005* | 0.556 |
|  | Stim*Electrode | 0.408 | 0.537 | 0.039 |
|  | Drug | 1.489 | 0.25 | 0.13 |
|  | Drug*Electrode | 1.296 | 0.281 | 0.115 |
|  | Stim*Drug | 0.079 | 0.784 | 0.008 |
|  | Stim*Drug*Electrode | 0.943 | 0.354 | 0.086 |
| Maximum net change in Hbo | Electrode | 1.805 | 0.209 | 0.153 |
|  | Stim | 15.061 | 0.003* | 0.601 |
|  | Stim*Electrode | 0.334 | 0.576 | 0.032 |
|  | Drug | 1.784 | 0.211 | 0.151 |
|  | Drug*Electrode | 1.06 | 0.327 | 0.096 |
|  | Stim*Drug | 0.114 | 0.743 | 0.011 |
|  | Stim*Drug*Electrode | 1.253 | 0.289 | 0.111 |
| Maximum net change in Hbr | Electrode | 1.334 | 0.275 | 0.118 |
|  | Stim | 19.48 | 0.001** | 0.661 |
|  | Stim*Electrode | 0.163 | 0.695 | 0.016 |
|  | Drug | 2.181 | 0.171 | 0.179 |
|  | Drug*Electrode | 0.749 | 0.407 | 0.07 |
|  | Stim*Drug | 0.234 | 0.639 | 0.023 |
|  | Stim*Drug*Electrode | 2.321 | 0.159 | 0.188 |

Surprisingly, in contrast to the significant effect on the nNOS IN-evoked haemodynamic response, changes in Hbt, Hbo and Hbr evoked by whisker stimulation were unchanged in the presence of LNAME (Fig 1F-J, Table 2-4). These data suggest that NO is not involved in the initiation of sensory-evoked functional hyperemia in the somatosensory cortex.

The emergence of an oscillation in all haemodynamic components on return to baseline following stimulation in the presence of LNAME was notable in both photostimulation- and whisker stimulation-evoked responses (Fig 1F), suggesting that NO may play a role in damping the haemodynamic return to baseline following either nNOS IN or whisker stimulation.

To confirm that the observed LNAME-associated differences in nNOS IN-evoked haemodynamic changes were not due to factors such as duration of anaesthesia, in a subset of mice experiments were also performed in the absence of pharmacological agents. No time-associated significant differences were observed in haemodynamic responses to either photostimulation or whisker stimulation, confirming that the described changes are not due to a timing effect (Fig 4, Table 5).

**Figure 4:**
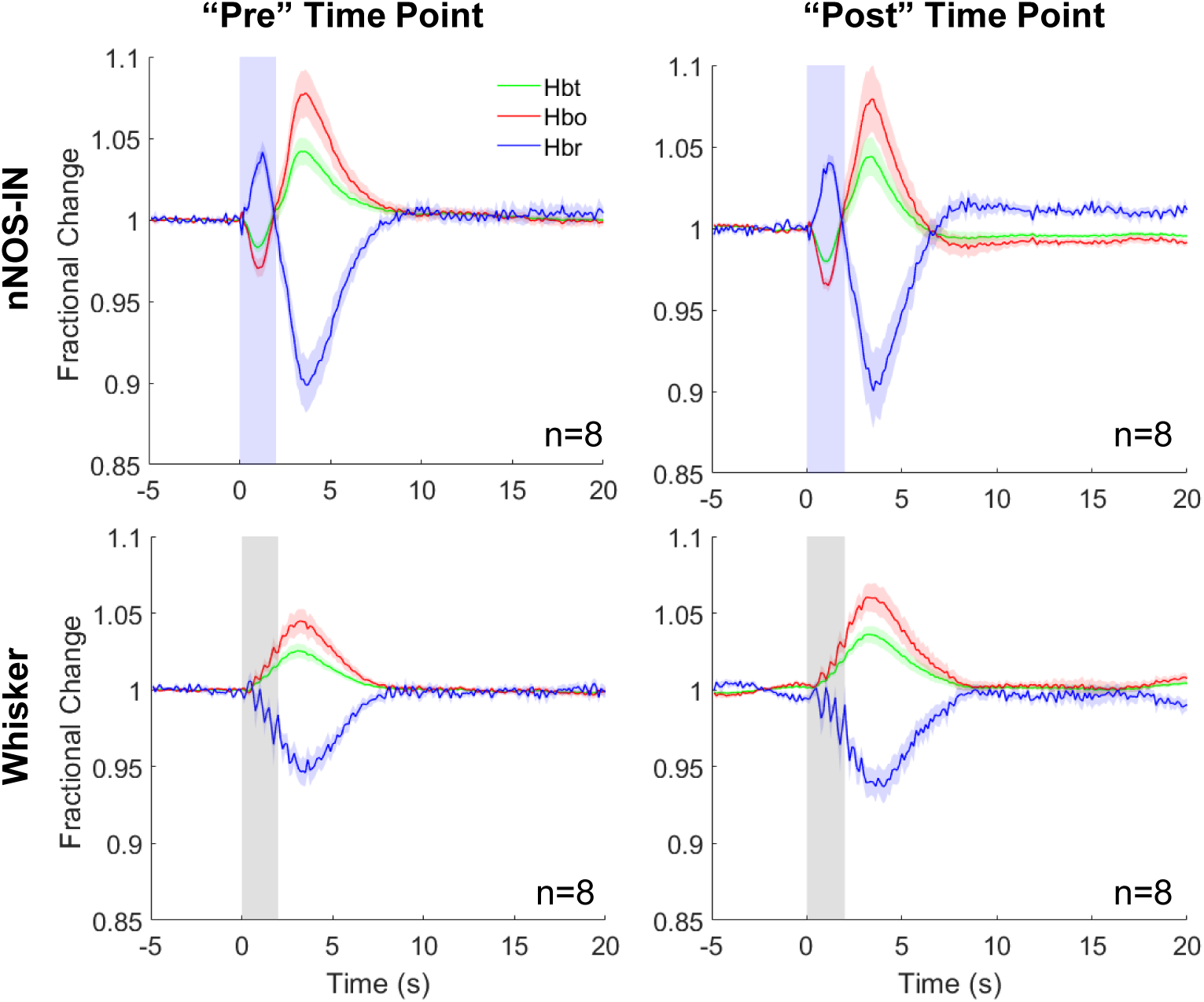
Time-matched experiments with no drug applied. Mean fractional change in Hbt, Hbo and Hbr in arteriolar ROI in response to 2s optogenetic activation of nNOS-INs (top row) or 2s whisker stimulation (bottom row) in experiments in which no pharmacological agent was applied. The “pre” (left) and “post” (right) measurement points were time-matched to the pre and post time points in the experiments in which LNAME was administered (Fig 1). Blue shading indicates photostimulation period (top row), grey shading indicates whisker stimulation period (bottom row). Data are mean ± SEM, n represents number of mice.

**Table 5:** Results of 2-way repeated measures ANOVA comparing haemodynamic responses in no drug condition (n=8 mice). *p<0.05, **p<0.01, ***p<0.001

| Variable | Factor | F(1,7) |  |  |
| --- | --- | --- | --- | --- |
| | | F | p | $\eta^2$ |
| Hbt response in initial dip period | Stim | 106.469 | 0.000017*** | 0.938 |
|  | Drug | 0.791 | 0.403 | 0.102 |
|  | Stim*Drug | 7.450 | 0.029 | 0.516 |
| Peak Hbt response | Stim | 2.895 | 0.133 | 0.293 |
|  | Drug | 1.946 | 0.206 | 0.218 |
|  | Stim*Drug | 1.133 | 0.322 | 0.139 |
| Maximum net change in Hbt | Stim | 14.348 | 0.007** | 0.672 |
|  | Drug | 1.866 | 0.214 | 0.21 |
|  | Stim*Drug | 0.113 | 0.747 | 0.016 |

Taken together, these data suggest that while NO plays a key role in the initial haemodynamic response evoked by nNOS IN activation, the initiation of whisker stimulation-evoked haemodynamic changes are largely independent of NOS activity.

### Evoked neural activity was unaltered by NOS inhibition

As the measured haemodynamic responses reflect stimulation-evoked neural activity, we assessed whether LNAME alters evoked neural activity. In a subset of mice, stimulation-evoked electrophysiological and haemodynamic changes were measured concurrently before and after LNAME injection to confirm that the observed LNAME effects reflect altered vascular responses, rather than a change in evoked neural activity. A sixteen channel NeuroNexus electrode was inserted into the centre of the whisker barrel cortex in order to measure the electrophysiological response to stimulation. In line with our previous findings (Lee et al., 2020), 2s photostimulation of cortical nNOS INs evoked a robust haemodynamic response (Fig 1) in the absence of a measurable change in ongoing neural activity (multi-unit activity (MUA); Fig. 3A). The lack of measurable change in neural activity persisted in the presence of LNAME (Fig 3A,C). Similarly, whisker stimulation-evoked increases in MUA, which extended throughout the depth of the cortex and lasted for the duration of the stimulation (Fig 3B), were unaffected by LNAME (Fig 3B-C). These data confirm that LNAME had no effect (Peak MUA: F(1,3)=0.032, p=0.869; Mean MUA: F(1,3)=0.003, p=0.958; Table 6) on the neural activity underlying the stimulation-evoked localised haemodynamic responses (Fig 1).

**Table 6:** Results of 2-way repeated measures ANOVA for stimulation-evoked MUA in absence and presence of LNAME (n=4 mice).

| Variable | Factor | F(1,3) |  |  |
| --- | --- | --- | --- | --- |
| | | F | p | $\eta^2$ |
| MUA Peak | Stim | 5.361 | 0.104 | 0.641 |
|  | Drug | 0.032 | 0.869 | 0.011 |
|  | Drug*Stim | 0.245 | 0.655 | 0.075 |
| Mean MUA | Stim | 3.865 | 0.144 | 0.563 |
|  | Drug | 0.003 | 0.958 | 0.001 |
|  | Drug*Stim | 0.007 | 0.939 | 0.002 |

### NO reduces 20-HETE-evoked vasoconstriction during nNOS IN activation

We hypothesised that the larger initial decrease in Hbt (indicative of vasoconstriction) observed in response to nNOS IN activation (Fig 1A,C,F,G) in the presence of the NOS inhibitor LNAME was due to 20-HETE production (Mulligan and MacVicar, 2004). To further investigate the NOS-dependent vasoactive pathway underlying nNOS IN-evoked localised haemodynamic changes, in a separate cohort of mice nNOS IN and whisker stimulation-evoked haemodynamic changes were recorded before and after (70-135mins) combined treatment with LNAME (75 mg/kg, i.p.) and HET0016 (N-(4-butyl-2-methylphenyl)-N’-hydroxy-methanimidamide, 10 mg/kg, i.v. (Poloyac et al., 2006), Fig 5), a selective inhibitor of CYP4A and CYP4F (Miyata et al., 2001). Repeated measures 2-way ANOVAs (factors: drug, stimulation type) revealed interactions between stimulation and drug for both the Hbt initial dip (F(1,5)=18.809, p=0.007, η^2^= 0.790) and peak (F(1,5)=23.137, p=0.005, η^2^= 0.822). Simple effects tests were therefore performed to assess the combined effect of LNAME and HET0016 on each stimulation type. For 2s photostimulation of nNOS INs, neither the initial dip in Hbt (Fig 5B, pre= 0.986±0.005, post= 0.973±0.01, p= 0.05), nor the peak Hbt (Fig 5C, pre= 1.07±0.011, post= 1.04±0.008, p= 0.052) were significantly different before and after treatment with LNAME and HET0016. Combined with the results of treating with LNAME alone (Fig 1), these data suggest that during short duration nNOS IN activation NO acts, at least in part, to reduce 20-HETE-elicited vasoconstriction.

**Figure 5:**
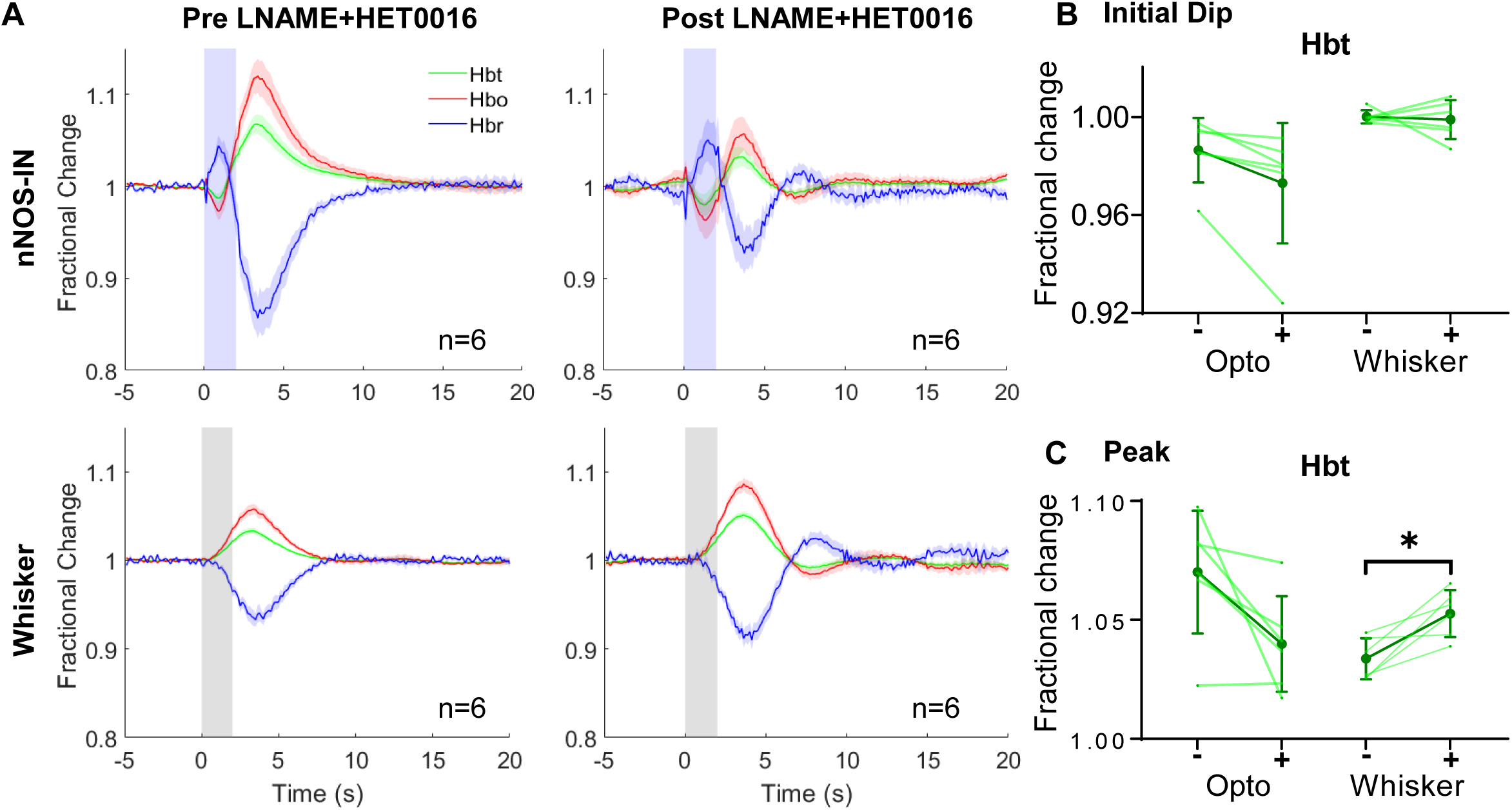
Haemodynamic responses evoked by nNOS-IN activation or whisker stimulation in presence of simultaneous inhibition of NOS and 20-HETE synthesis. Group data (n=6 mice). **(A):** Mean fractional change in Hbt, Hbo and Hbr in arteriolar ROI before (left) and after (right) LNAME and HET0016 injection. Blue shading indicates photostimulation period (upper panels), grey shading indicates whisker stimulation period (lower panels). Data are mean ± SEM. **(B-C):** Darker lines represent group mean ± SD, lighter lines indicate trial-averaged mean for individual animals. **(B):** Initial fractional change (‘initial dip’) in Hbt in response to optogenetic and whisker stimulation, with (+) and without (-) inhibitors (LNAME+HET0016). **(C):** Maximum fractional change in Hbt (‘peak’) evoked by optogenetic and whisker stimulation, with (+) and without (-) LNAME+HET0016. *p<0.05

In response to 2s whisker stimulation, the peak Hbt change was significantly greater when production of NO and 20-HETE were inhibited (Fig 5C, pre= 1.034±0.004, post= 1.053±0.004, p= 0.013), suggesting that production of 20-HETE during sensory stimulation causes a constriction which is overcome by an NO-independent vasodilatory mechanism.

As previously observed in the presence of LNAME alone (Fig 1F), an oscillation in all haemodynamic components was observed on return to baseline following stimulation in the presence of LNAME and HET0016 (Fig 5A), further supporting our suggestion that NO plays a role in damping the haemodynamic return to baseline following nNOS IN activation or sensory stimulation.

### Haemodynamic responses to whisker stimulation and nNOS IN activation sum in a linear manner

Having demonstrated the differential involvement of NO and 20-HETE in the haemodynamic responses to nNOS IN activation and whisker stimulation, we hypothesised that whisker stimulation-evoked functional hyperemia and nNOS IN- evoked haemodynamic responses are driven by different vasoactive pathways. If this is the case, summing the change in Hbt evoked by photostimulation of cortical nNOS INs and that evoked by whisker stimulation should predict the change in Hbt evoked by simultaneous presentation of whisker and nNOS IN stimulation. To test this, a subset of animals received simultaneous photostimulation of nNOS INs and whisker stimulation, in addition to the separate photostimulation and whisker stimulation described above. In these mice, linear summation of the Hbt time series evoked by whisker stimulation and that evoked by photostimulation of cortical nNOS INs predicted the time series of the Hbt changes evoked by simultaneous stimulation of nNOS INs and whiskers, both in the absence and presence of LNAME (Fig 6). This linear summation further supports our suggestion that nNOS IN activation and whisker stimulation drive haemodynamic changes via different vasoactive pathways.

**Figure 6:**
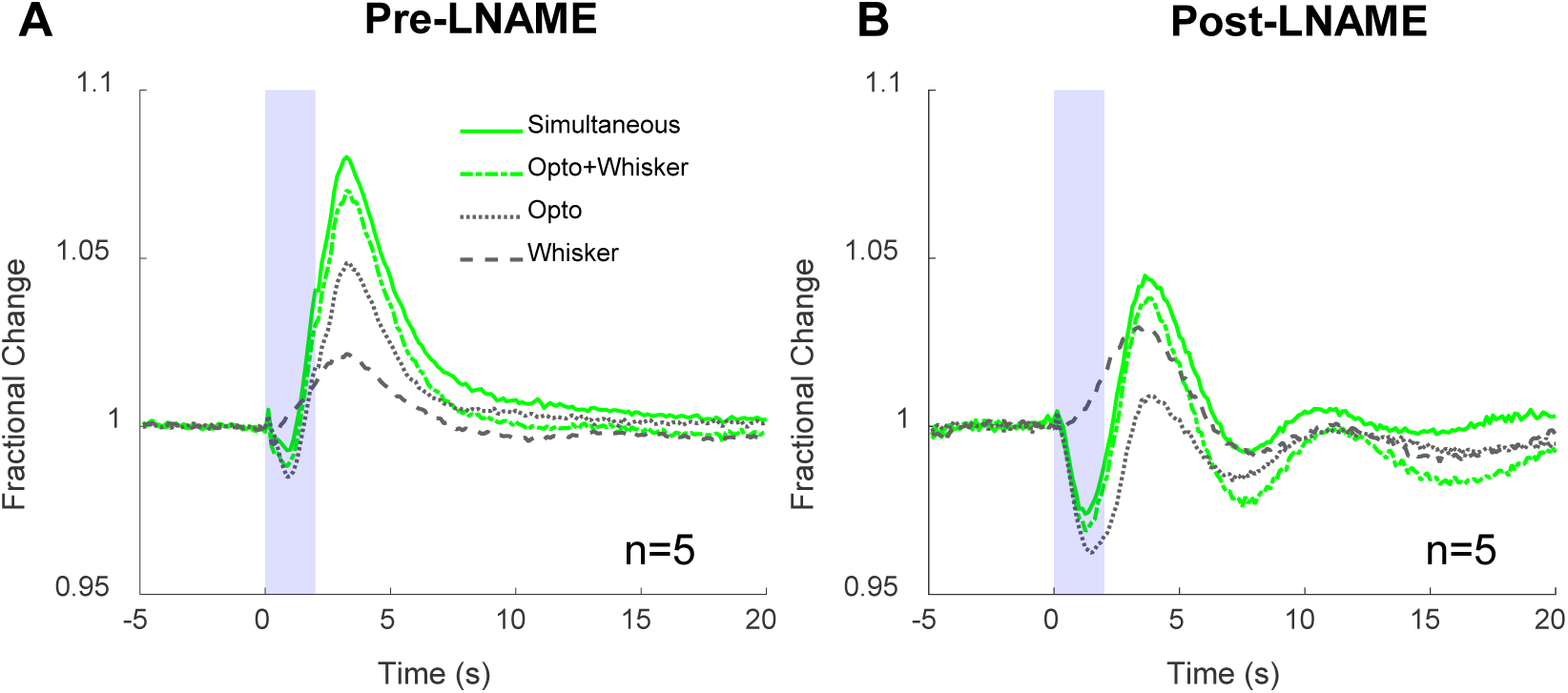
Change in Hbt evoked by simultaneous presentation of whisker and optogenetic stimulation is similar to that predicted by summing changes in Hbt evoked by independent optogenetic and whisker stimulation. Group data (n=5 mice): Mean fractional change in Hbt in arteriolar ROI before **(A)** and after **(B)** NOS inhibition with LNAME. Blue shading indicates stimulation period. Continuous green line indicates haemodynamic response to 2s simultaneous stimulation, dash and dot green line indicates the predicted response calculated by summing the responses to separate 2s optogenetic (grey dotted line) and 2s whisker (grey dashed line) stimulations. For visual clarity, error bars are not shown. (Separate optogenetic and whisker stimulation data are also included in Figure 1).

### Systemic injection of LNAME enhances vasomotion in anaesthetised mice

NO can suppress vasomotion (Dirnagl et al., 1993a), a low frequency oscillation in arteriole diameter occurring at ∼0.1 Hz (Drew et al., 2011; Mayhew et al., 1996). Therefore, in addition to characterising the effect of NOS inhibition by LNAME on stimulation-evoked cortical haemodynamics, we also examined the effect of NOS inhibition on low frequency arterial oscillations. We detected an increase in the power of low frequency arterial oscillations, centred around 0.1Hz, after LNAME injection as compared to before injection (Fig 7, area under the curve (AUC): [0.09-0.11 Hz]; p=0.000062, Table 7-8), which was not apparent in the absence of LNAME (Fig 7, ‘pre’ vs ‘post’ in no drug condition, p=0.531, Table 7-8). These data confirm that NOS inhibition results in enhanced low frequency vascular oscillations (Ances et al., 2010; Biswal and Hudetz, 1996). Additional peaks in the power spectrum which reflect the frequency of stimulation (ISI of 25s: 0.04Hz) and its harmonics can be seen in all cases (Fig 7). As stimulation paradigms were interleaved and LNAME alters the haemodynamic response to photostimulation of nNOS INs but not whisker stimulation (Fig 1), the peak associated with the stimulation pattern is shifted to a lower frequency in the presence of LNAME (Fig 7B).

**Figure 7:**
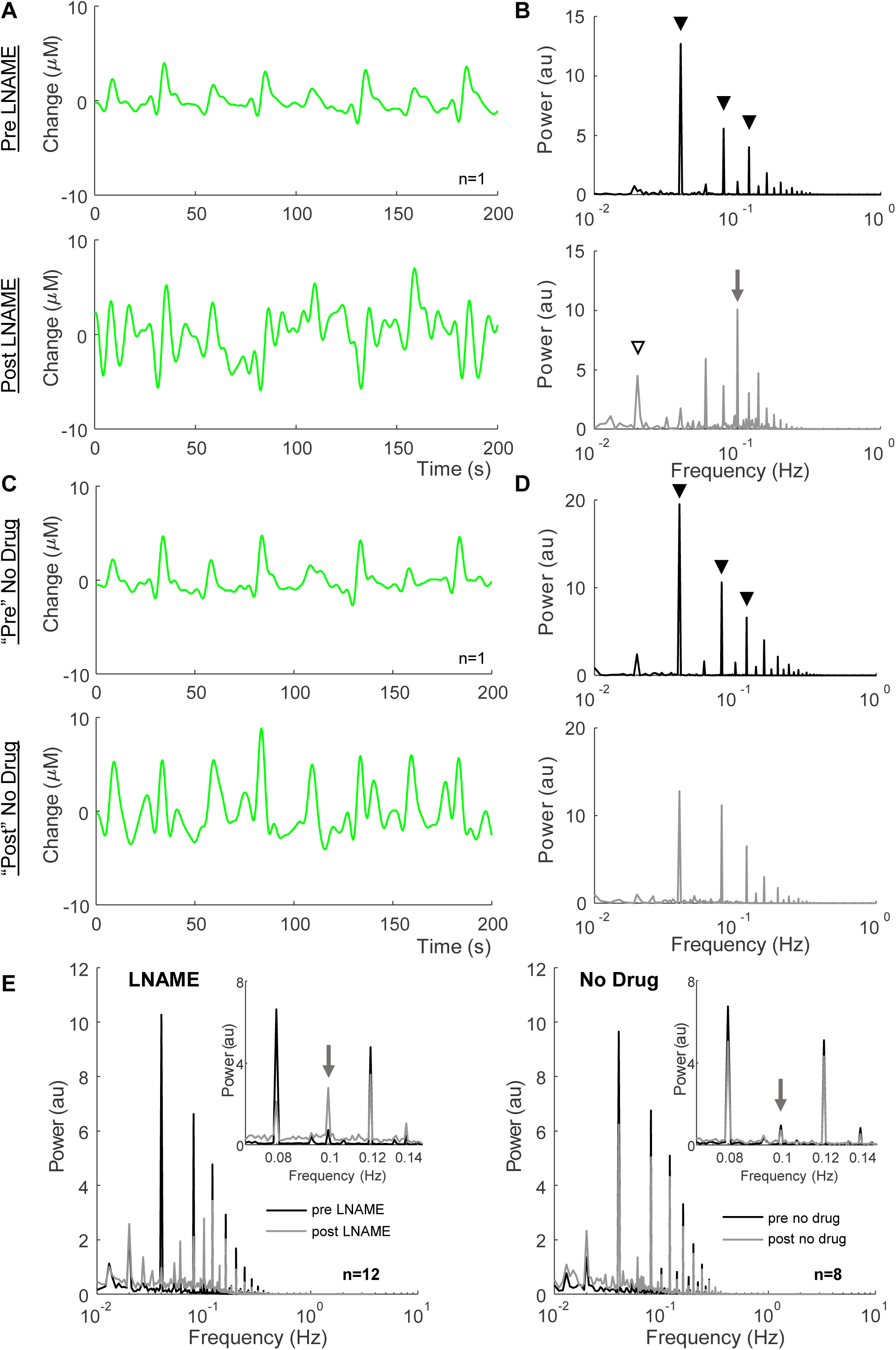
LNAME enhances vasomotion. **A,C)**: Example 200s Hbt time series from experiments occurring before (upper) and after (lower) LNAME injection **(A)** or no drug treatment **(C)**. Responses to individual stimulations can be seen. **(B,D):** Example power spectrum of Hbt data (1000s duration analysed) from same experiments as A,C, before (upper, black) and after (lower, grey) LNAME injection **(B)** or no drug treatment **(D)**. Peaks are observed at frequency of stimulation (ISI 25s: 0.04Hz) and its harmonics (black arrowheads). After LNAME injection, a peak at 0.1Hz (vasomotion, grey arrow) is observed and the apparent stimulation frequency is reduced (white arrowhead) **(E):** Mean power spectrum of oscillations in Hbt before (black) and after (grey) LNAME (left, n=12) or no drug treatment (right, n=8). **Inset**: highlight of 0.07-0.15Hz. Data from arteriolar ROI in experiments as shown in Figures 1 and 4. For visual clarity, error bars are not shown. n indicates number of mice.

**Table 7:**
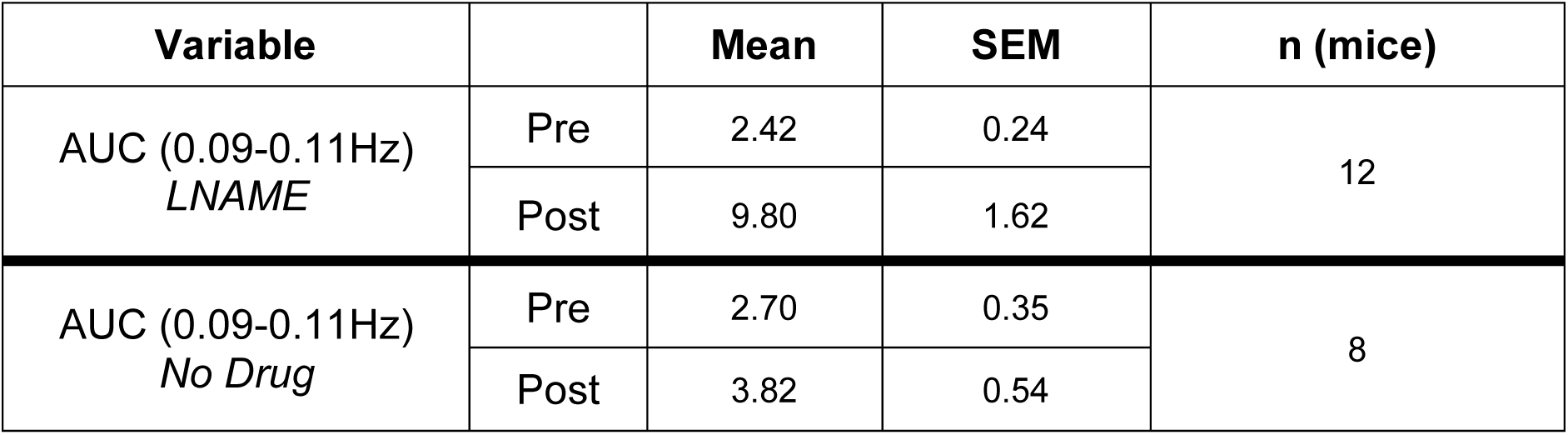
Descriptive statistics of power in low frequency arteriole oscillations (AUC, 0.09-0.11Hz).

| Variable |  | Mean | SEM | n (mice) |
| --- | --- | --- | --- | --- |
| AUC (0.09-0.11Hz)<br>*LNAME* | Pre | 2.42 | 0.24 | 12 |
|  | Post | 9.80 | 1.62 |  |
| AUC (0.09-0.11Hz)<br>*No Drug* | Pre | 2.70 | 0.35 | 8 |
|  | Post | 3.82 | 0.54 |  |

**Table 8:** Statistical analysis of power in low frequency arteriole oscillations: 2 way mixed ANOVA (*p<0.05, **p<0.01, ***p<0.001) and simple effects tests (following Bonferroni correction for multiple comparisons, *p<0.0125, **p<0.0025, ***p<0.00025).

| Variable | Factor | F(1,18) |  |  |
| --- | --- | --- | --- | --- |
| | | F | p | $\eta^2$ |
| AUC (0.09-0.11Hz) | Time | 14.265 | 0.001** | 0.442 |
|  | Drug | 8.948 | 0.008** | 0.332 |
|  | Drug*Time | 7.765 | 0.012* | 0.301 |

| Simple Effects Tests: time |  | p |
| --- | --- | --- |
| LNAME | Post vs Pre | 0.000062*** |
| No Drug | Post vs Pre | 0.531 |
| Simple Effects Tests: drug |  | p |
| Pre timepoint | LNAME vs No Drug | 0.497 |
| Post timepoint | LNAME vs No Drug | 0.009* |

## Discussion

Combining *in vivo* 2D-OIS, electrophysiology, optogenetics and pharmacology, we investigated the role of NO in nNOS IN driven and sensory-evoked neurovascular coupling in the somatosensory cortex of mice. We demonstrated that: first, localised nNOS IN driven haemodynamic changes are mediated by NO; second, initiation of the haemodynamic response to short duration sensory stimulation is not dependent on NO; and third, NO suppresses vasomotion in anaesthetised mice.

By inhibiting NO synthesis with LNAME, we demonstrated the key role that NO plays in the haemodynamic response to nNOS IN activation (Fig 1). Inhibiting production of NO by NOS revealed a larger initial decrease in Hbt in response to nNOS IN activation, which was attenuated when production of NO and 20-HETE were inhibited simultaneously. However, following initiation of the haemodynamic response, the successive increase in Hbt was unaltered by NOS inhibition. These data suggest that the initial vasodilatory response elicited by nNOS IN activation is mediated by NO, at least in part by inhibition of constriction by 20-HETE, while a later vasodilation involves an NO-independent mechanism. These findings extend previous work in cortical slices suggesting that type II nNOS INs evoke vasodilation by NO release (Perrenoud et al., 2012). A likely candidate for the delayed vasodilation is GABA (Fergus and Lee, 1997), which is released by active nNOS INs and may act via astrocytes to elicit haemodynamic changes (Cauli and Hamel, 2010; Lecrux and Hamel, 2016).

Our demonstration of an nNOS IN-elicited vasoconstriction (which was increased when NOS was inhibited) further supports the idea that arteriolar constrictions may be evoked by INs, rather than by excitatory neurons (Uhlirova et al., 2016). The observed increase in Hbr elicited by nNOS IN activation (Fig 1F) is analogous to the negative BOLD signal, further linking IN activity to negative BOLD fMRI responses (Boorman et al., 2015, 2010; Lee et al., 2020).

Whilst NO was found to mediate nNOS IN driven changes in cerebral blood volume, the neurovascular coupling response to short duration sensory stimulation was largely unaltered when NO production was reduced (Fig 1). A lack of effect on stimulation-evoked neural activity (Fig 3) confirmed that sensory-evoked neurovascular coupling was also unaffected by treatment with LNAME. Although nNOS- and eNOS-derived NO have been suggested to play a key role in sensory-evoked neurovascular coupling (Ayata et al., 1996; Chow et al., 2020; Dirnagl et al., 1993b; Hosford and Gourine, 2019; Toth et al., 2015) our results call into question both the involvement of nNOS INs in sensory-evoked functional hyperemia (Echagarruga et al., 2020) and that NO release is an important mediator of sensory-evoked haemodynamic responses (Hosford and Gourine, 2019). Our findings are supported by previous studies which have also reported a limited effect of NOS inhibition (Adachi et al., 1994; Lindauer et al., 1999; Vazquez et al., 2018) or genetic deletion of nNOS (Liu et al., 2008; Ma et al., 1996) on sensory-evoked functional hyperemia.

Discrepancies in the literature may be due to differences in experimental approaches. First, the effect of LNAME may be affected by the application route chosen. Here we used i.p. injection of LNAME to reduce NOS activity throughout the brain (Bannerman et al., 1994; Salter et al., 1995), as compared to topical application which may result in effects being restricted to the superficial cortical layers, or intracortical injection which may limit the effect on pial arterioles (Vazquez et al., 2018) - the vascular component most affected by NO (Echagarruga et al., 2020; Mishra et al., 2016). Second, stimulation duration may dictate the effect of NOS inhibition on the evoked haemodynamic response, with opposing effects being previously reported for long (60s) and short (4s) duration sensory stimulation (Ances et al., 2010). Therefore, the discrepancy between our results and those of Liu et al. (2008) who used a similar pharmacological approach to report a significant involvement of NO (via inhibition of 20-HETE production) in the haemodynamic response to sensory stimulation in rats, may be at least partly explained by the use of short (2s, here) and long (60s, (Liu et al., 2008)) duration stimulations.

While we found no evidence for the involvement of NO in the initiation of sensory-evoked functional hyperemia, following inhibition of NO production we observed an oscillation in Hbt, Hbo and Hbr on return to baseline at stimulation offset in both the photostimulation and whisker stimulation experiments (Fig 1F). The shape of the haemodynamic response indicates that the haemodynamic return to baseline following stimulation is underdamped, suggesting that NO acts to dampen vasoconstriction after stimulation ends (Dormanns et al., 2016; Kenny et al., 2018). A similar arteriolar constriction, sometimes seen following sensory stimulation, has been shown to be mediated by NPY (Uhlirova et al., 2016). Future work could elucidate whether NO attenuates the vasoconstrictive actions of NPY release from nNOS INs following sensory stimulation.

In agreement with previous studies (Ances et al., 2010; Biswal and Hudetz, 1996; Dirnagl et al., 1993a) we demonstrated a role for NO in suppressing vasomotion. When NO production was attenuated with LNAME, we observed an increase in the power of low frequency haemodynamic oscillations centred at ∼0.1Hz, compared to before injection of LNAME (Fig. 7). LNAME may lead to enhanced vasomotion via the inhibition of eNOS-derived NO, which has previously been suggested to inhibit voltage gated calcium channels in arterial smooth muscle cells and, thereby, vasomotion (Yuill et al., 2010).

Our study is not without limitations. Due to the use of a non-specific NOS inhibitor, it is not possible for us to dissect the involvement of eNOS and nNOS-associated vasoactive mechanisms in the vascular responses we observed. As both the cellular specificity of NOS isoform expression and the isoform-specificity of pharmacological agents have recently been questioned (Echagarruga et al., 2020), designing experiments that allow specific cellular or isoform dissection is difficult.

2D-OIS, used in this study to investigate cortical haemodynamics, predominantly reflects changes in haemoglobin concentration in superficial blood vessels. Although previous studies have suggested that NO is preferentially involved in the regulation of pial arterial, as opposed to capillary, diameter (Echagarruga et al., 2020; Mishra et al., 2016), further studies using approaches such as 2-photon microscopy would allow assessment of whether nNOS INs can evoke capillary dilation.

Anaesthesia allows the involvement of NO in the regulation of CBF to be assessed in the absence of behaviours such as locomotion, which has been shown to evoke NO-dependent vasodilation (Echagarruga et al., 2020) and can significantly alter sensory stimulus-evoked haemodynamic responses (Eyre et al., 2022). Furthermore, the prolonged temporal nature of the experimental paradigm used in this study makes it difficult to perform in awake animals. However, anaesthesia may confound neurovascular coupling (Gao et al., 2017) and may also affect the recruitment of INs to sensory stimulation (Vazquez et al., 2018), thereby potentially altering the reliance of functional hyperemia on NO. To minimise the confound of anaesthesia we used an anaesthetic regime under which haemodynamic responses are similar to those observed in the awake mouse (Sharp et al., 2015).

Understanding how, and when, nNOS INs, and NO, regulate CBF may highlight novel therapeutic strategies for neurodegenerative diseases. CBF deficits are observed early in AD (Iturria-Medina et al., 2016) and are linked to cognitive decline (Hays et al., 2016; Leijenaar et al., 2017). Therefore, targeting nNOS INs, or increasing NO bioavailability (Lourenço and Laranjinha, 2021), to reverse CBF deficits may prove to be a beneficial strategy for preventing cognitive decline.

## Materials and Methods

### Animals

All procedures involving animals were performed in accordance with the UK Government, Animals (Scientific Procedures) Act 1986, were approved by the University of Sheffield ethical review and licensing committee, and were in line with the ARRIVE guidelines (Percie du Sert et al., 2020). Mice had ad libitum access to food and water and were housed on a 12 hour dark/light cycle. 23 mice were used (14 males and 9 females), aged 4-9 months old (6.58 ± 1.36months) and weighing 20-36g (29.08 ± 5.73g) on date of surgery. Mice were nNOS-CreER x ChR2-EYFP (nNOS-ChR2), obtained by crossing heterozygous nNOS-CreERT mice (Stock 014541, Jackson Laboratory (Taniguchi et al., 2011)) with homozygous ChR2(H134R)-EYFP mice (Stock 024109, Jackson Laboratory (Madisen et al., 2012)), as used previously (Lee et al., 2020). Mice positive for the nNOS-CreERT insertion (confirmed by genotyping of ear clips from pups) were used in these experiments. ChR2 expression was induced by intraperitoneal (i.p.) injection of tamoxifen (Sigma-Aldrich) at 100mg/kg, administered 3 times with a day between each injection. Treatment with tamoxifen was carried out when mice were aged between 1-5 months, and took place a minimum of 2 weeks prior to surgery to allow for gene expression to occur. Randomisation sequences were not used to assign animals to different pharmacological agents.

### Surgical Preparation of Chronic Cranial Window

At least 2 weeks prior to experimental sessions, surgery to prepare a thinned cranial window over the right somatosensory cortex was performed as previously described by Sharp et al. (2015). In brief, i.p. injection of fentanyl-fluanisone (Hypnorm, Vetapharm Ltd), midazolam (Hypnovel, Roche Ltd) and sterile water (in a ratio of 1:1:2 by volume; 7ml/kg) was used to induce anaesthesia and anaesthesia was maintained using isoflurane (0.5-0.8%) in 100% oxygen at a flow rate of 1L/min. Surgical anaesthetic plane was monitored by regularly checking toe-pinch reflex response. To avoid optogenetic activation of ChR2, surgeries were performed in a dark room using a surgical light with a band pass filter (577±5 nm). Mice were placed in a stereotaxic frame (Kopf Instruments) with a homoeothermic blanket (Harvard Apparatus) maintaining rectal temperature at 37°C. Using a dental drill, an area of bone (∼3mm x 3mm) overlying the right whisker barrel cortex was thinned to translucency and a thin layer of clear cyanoacrylate was applied to smooth and reinforce the area. Dental cement (Super bond C&B, Sun Medical) was used around the window to secure a stainless steel head-plate to the skull. Following surgery, mice were singly housed and were monitored using the mouse grimace scale (Langford et al., 2010) and weighed weekly. Any animals losing over 20% body weight post-operatively were culled; for this study no mice met this criteria.

### 2-Dimensional Optical Imaging Spectroscopy

Anaesthesia was induced via i.p. injection of fentanyl-fluanisone (Hypnorm, Vetapharm Ltd), midazolam (Hypnovel, Roche Ltd) and sterile water (in a ratio of 1:1:2 by volume; 7ml/kg) and anaesthesia was maintained using isofluorane (0.25%-0.7%) with 100% oxygen at a flow rate of 0.8L/min. Mice were head fixed on a stereotaxic frame, using the head-plate that was attached during surgery. A homeothermic blanket maintained rectal temperature of the mouse at 37°C. 2-Dimensional Optical Imaging Spectroscopy (2D-OIS) allows changes in cortical haemodynamics (oxygenated haemoglobin: Hbo, deoxygenated haemoglobin: Hbr and total haemoglobin: Hbt) to be measured. 2D-OIS was performed using our previously published methodology (Berwick et al., 2005). In brief, using a Lambda DG-4 high-speed filter changer (Sutter Instrument Company, USA), the cortex under the thinned window was illuminated with four wavelengths of light (587±9nm, 595±5nm, 560±15nm, 575±5nm). The remitted light was collected with a Dalso 1M60 CCD camera with a frame rate of 32 Hz. This was synchronised to the wavelength switching, resulting in an effective frame rate of 8Hz. To avoid collecting light from the 470nm photostimulation LED, the camera was fitted with a 490nm high pass filter. Spectral analysis was carried out pixel-by-pixel on the remitted light, based on the path length scaling algorithm (PLSA) as described previously (Berwick et al., 2005; Mayhew et al., 1999), which allowed the generation of estimates for the changes from baseline of Hbo, Hbr and total haemoglobin Hbt. The analysis assumed a baseline tissue concentration of haemoglobin of 100μM and oxygen saturation of 80%. This spectral analysis resulted in the production of 2-dimensional spatial images representing micromolar changes in the concentration of Hbo, Hbr and Hbt, over the time course of an experiment.

### Electrophysiology

In some experiments neural activity was recorded concurrently with haemodynamic activity, to effectively assess both these aspects of neurovascular coupling. A cranial burr-hole was made in the right whisker barrel cortex (identified by response to whisker stimulation in a previous 2D-OIS experiments) to allow insertion of a 16 channel microelectrode (100μm spacing, 1.5-2.7Ω impedance, site area 177μm^2^; Neuronexus Technologies) to a depth of ∼1600μm. To record neural activity, the electrode was connected to a preamplifier and data acquisition device (Medusa BioAmp/ RZ5, TDT), and data were collected at 24kHz. During post-hoc analysis, data were downsampled to 6kHz and a 500Hz high pass filter was applied. A ‘spike’ was detected when data exceeded a threshold of 1.5 times the standard deviation above the mean. The number of spikes occurring in 100ms bins were counted and reported as MUA. Data from the 12 electrode channels corresponding to cortical depth were used for analysis. To produce fractional change in MUA, responses were normalised to a 2s baseline. Trials were averaged to produce a mean response for each stimulation paradigm for each animal. For statistical analysis, the peak MUA and mean MUA during the stimulation period were calculated for each animal. Group means were produced by averaging across animals. One mouse was excluded from statistical analysis due to the presence of excessive noise.

### Stimulations

#### Photostimulation

A fibre-coupled 470nm LED light source (ThorLabs) connected to a fibre optic cable (core diameter 200μm, Thorlabs, USA) delivered blue light for photostimulation of ChR2. 2s light delivery consisted of 10ms pulses at 99Hz (1V, 0.45 mW).

#### Whisker stimulation

A plastic T-bar attached to a stepper motor moved whiskers of the left whisker pad ∼1cm in the rostro-caudal direction (Sharp et al., 2015). Whiskers were stimulated for 2s, at 5Hz.

*Simultaneous presentation* of photostimulation and whisker stimulation was applied in order to assess whether the evoked haemodynamic responses summed in a linear manner.

In order to mitigate the effects of time since anaesthesia or pharmacological treatment, optogenetic, whisker and simultaneous stimulations were interleaved within each experiment. To improve the signal to noise ratio, each experiment contained 20-30 repeats of each stimulation type, with an inter-stimulus interval (ISI) of 25s.

### Pharmacology

To reduce NO production, mice were treated with the non-selective NOS inhibitor N(G)-Nitro-L-arginine methyl ester (LNAME, Sigma). LNAME (10mg/ml made up with sterile saline) was administered via i.p. bolus injection (75mg/kg; which has been shown to reduce NOS activity within the cortex by 93% within 1 hour of injection (Bannerman et al., 1994)). 20-HETE production was inhibited by treatment with a selective inhibitor of CYP4A and 4F, N-(4-butyl-2-methylphenyl)-N’-hydroxy-methanimidamide (HET0016: Miyata et al., 2001). HET0016 (Santa Cruz Biotechnology) was administered i.v. (tail vein, 10 mg/kg; a 5mg/ml solution was made up with 10% lecithin saline (Poloyac et al., 2006; Singh et al., 2007)).

### Procedure

Following anaesthetic induction, an electrode was inserted into the right whisker barrel cortex (if needed). 2D-OIS measurements of haemodynamic changes evoked by stimulation (photostimulation and/or whisker stimulation) were monitored continuously. Injection of agent(s) occurred approximately 1 hour after 2D-OIS experiments started which, in the case of electrode insertion, allowed sufficient time for haemodynamics to recover from the resulting CSD (Chang et al., 2010). Timings of agent injection were kept consistent, regardless of whether an electrode was inserted. For this study, data from specific timepoints pre- and post-treatment were analysed. Pre-treatment responses to stimulation were taken from the experiment immediately prior to the injection of pharmacological agent(s). Responses to stimulation post-treatment were taken from the experiment occurring at the following time after injection of pharmacological agent(s): LNAME: 70-135 minutes (mean= 108±7 minutes) after injection which, in agreement with previous studies (Bannerman et al., 1994; Salter et al., 1995), was sufficient time for LNAME to have maximal effect (Fig 2); LNAME and HET0016: 70-135 minutes (121±10 minutes) after injection; No drug (timing control experiments): 80-120 minutes (100±6 minutes) after the ‘injection’ timepoint.

### Data analysis

All experiments and analysis were performed unblinded. Data analysis was performed using MATLAB (MathWorks). Using the spatial map of Hbt changes evoked by 2s whisker stimulation, generated by 2D-OIS, a region of interest (ROI) was automatically selected (Lee et al., 2020). In brief, each pixel was averaged across time during the response period, to generate a mean pixel value. Any pixel whose value was greater than 1.5x standard deviation was included in the ROI. The resulting ROI (white ROI, Fig 1B) represents the area with the greatest haemodynamic response to the whisker stimulation, and therefore represents the whisker barrel cortex. Within this ROI, the arterial region most responsive to the pharmacological intervention was manually selected (red ROI, Fig 1B). For analysis of responses within the arterial ROI, baseline oxygen saturation and tissue concentration of haemoglobin of 80% and 100μM, respectively, were assumed. For post-treatment data, baseline assumptions were corrected based on the measured change in Hbt and Hbo evoked by treatment with the pharmacological agent(s). For each stimulation paradigm, the response across all pixels within the arterial ROI was averaged, producing the three haemodynamic time series (Hbo, Hbr, Hbt).

#### Comparison of stimulation-evoked responses

To compare stimulation-evoked responses in trials involving photostimulation, residual artefact from the 470nm LED was removed using a modified boxcar function. Micromolar changes in haemodynamics were converted to fractional change (as compared to a 5s baseline). For each stimulation paradigm, mean time series were produced for each animal by averaging across trials. To produce the group mean time series, these responses were then averaged across animals within each group.

#### Low frequency vascular oscillations

The last 1000s of the Hbt timeseries of ‘pre-’ and ‘post-’ experiments were used to assess low frequency vascular oscillations. A cubic polynomial trend was removed from the data and the amplitude of the signal was normalised between 0 and 1, before computing the Fourier transform (FFT). The area under the curve (AUC) between 0.09-0.11Hz was calculated in order to compare oscillations centred around 0.1Hz (vasomotion). To produce the group mean, FFTs (and associated AUC) were averaged across animals within each group.

For visualisation purposes (Fig 7A-C), following removal of a cubic polynomial trend a filter was applied to the individual Hbt time series to remove high frequency components (such as heart rate).

### Statistical analysis

Three metrics (Fig 1G) were extracted from the stimulation-evoked haemodynamic time series: *initial dip*: the minimum value of Hbt and Hbo and maximum value of Hbr during the initial response period (0.25-5s after stimulation onset), *peak:* the maximum value of Hbt and Hbo and minimum value of Hbr during the response period (0.25-10s after stimulation onset) and *net change:* the magnitude of net change in Hbt, Hbo or Hbr (maximum - minimum value, above). Statistical analysis was carried out using SPSS (version 26). Shapiro-Wilk test was used to assess normality of data (ANOVA was considered robust if data were approximately normally distributed) and Levene’s test was used to test for equality of variances. Outliers were identified as extreme if they had a studentised residual >3. ANOVAs are considered robust against minor violations of assumptions. For experiments in which LNAME was applied alone, to determine statistical significance, a 3-way mixed ANOVA (within-group factors: drug [pre/post], stimulation type [photostimulation/whisker]; between-group factor: electrode [absent/present]) was used. For other pharmacological agents and MUA, a repeated measures 2-way ANOVA was used (factors: drug [pre/post], stimulation type [photostimulation/whisker]). To assess differences in arterial oscillations, spectral power (AUC) was compared using a mixed 2-way ANOVA (between-group factor: drug [LNAME/no drug]; within-group factor: time [pre/post agent application]). In the spectral power comparison, one data point was identified as an extreme outlier, however its inclusion did not alter the outcome of the statistical analysis and so all data were included. For all multi-way ANOVAs, Greenhouse-Geisser correction was applied and simple effects tests were carried out to further interrogate any significant interaction effects. To assess the evolution of the effect of LNAME, a 1-way ANOVA was performed, comparing 4 timepoints (pre-LNAME injection, 0 minutes after LNAME injection, 60-70 minutes after, and 95-135 minutes after). Results were considered statistically significant if p < 0.05, unless otherwise stated. All data are reported as mean ± standard error of the mean (SEM), unless otherwise stated. Sample sizes were based on those in previously published studies using similar pharmacological approaches (Bannerman et al., 1994).

### Data availability

Data files used/analysed in the current study are available in the DRYAD repository.

## Acknowledgements

CH was funded by a Sir Henry Dale Fellowship jointly funded by the Wellcome Trust and the Royal Society. This research was funded in whole, or in part, by the Wellcome Trust [Grant number 105586/Z/14/Z]. For the purpose of Open Access, the author has applied a CC BY public copyright licence to any Author Accepted Manuscript version arising from this submission. This work was funded in part by the Medical Research Council UK [Grant number MR/M013553/1 to J.B and L.B]. We would like to thank Michael Port for building and maintaining the whisker stimulation device and 2D-OIS imaging equipment, and Rachel Sandy for her assistance with tail vein injections.

## References

Adachi, K., Takahashi, S., Melzer, P., Campos, K.L., Nelson, T., Kennedy, C., Sokoloff, L., 1994. Increases in local cerebral blood flow associated with somatosensory activation are not mediated by NO. Am J Physiol 267, H2155–2162. https://doi.org/10.1152/ajpheart.1994.267.6.H2155

Alonso-Galicia, M., Hudetz, A.G., Shen, H., Harder, D.R., Roman, R.J., 1999. Contribution of 20-HETE to vasodilator actions of nitric oxide in the cerebral microcirculation. Stroke 30, 2727–2734; discussion 2734. https://doi.org/10.1161/01.str.30.12.2727

Ances, B.M., Greenberg, Joel. H., Detre, J.A., 2010. Interaction between Nitric Oxide Synthase Inhibitor Induced Oscillations and the Activation Flow Coupling Response. Brain Res 1309, 19–28. https://doi.org/10.1016/j.brainres.2009.09.119

Anenberg, E., Chan, A.W., Xie, Y., LeDue, J.M., Murphy, T.H., 2015. Optogenetic stimulation of GABA neurons can decrease local neuronal activity while increasing cortical blood flow. J. Cereb. Blood Flow Metab. 35, 1579–1586. https://doi.org/10.1038/jcbfm.2015.140

Attwell, D., Buchan, A.M., Charpak, S., Lauritzen, M., MacVicar, B.A., Newman, E.A., 2010. Glial and neuronal control of brain blood flow. Nature 468, 232–243. https://doi.org/10.1038/nature09613

Ayata, C., Ma, J., Meng, W., Huang, P., Moskowitz, M.A., 1996. L-NA-Sensitive rCBF Augmentation during Vibrissal Stimulation in Type III Nitric Oxide Synthase Mutant Mice. J Cereb Blood Flow Metab 16, 539–541. https://doi.org/10.1097/00004647-199607000-00002

Bannerman, D.M., Chapman, P.F., Kelly, P.A., Butcher, S.P., Morris, R.G., 1994. Inhibition of nitric oxide synthase does not prevent the induction of long-term potentiation in vivo. J Neurosci 14, 7415–7425.

Beishon, L., Clough, R.H., Kadicheeni, M., Chithiramohan, T., Panerai, R.B., Haunton, V.J., Minhas, J.S., Robinson, T.G., 2021. Vascular and haemodynamic issues of brain ageing. Pflugers Arch 473, 735–751. https://doi.org/10.1007/s00424-020-02508-9

Berwick, J., Johnston, D., Jones, M., Martindale, J., Redgrave, P., McLoughlin, N., Schiessl, I., Mayhew, J.E.W., 2005. Neurovascular coupling investigated with two-dimensional optical imaging spectroscopy in rat whisker barrel cortex. European Journal of Neuroscience 22, 1655–1666. https://doi.org/10.1111/j.1460-9568.2005.04347.x

Biswal, B.B., Hudetz, A.G., 1996. Synchronous oscillations in cerebrocortical capillary red blood cell velocity after nitric oxide synthase inhibition. Microvasc Res 52, 1–12. https://doi.org/10.1006/mvre.1996.0039

Boorman, L., Harris, S., Bruyns-Haylett, M., Kennerley, A., Zheng, Y., Martin, C., Jones, M., Redgrave, P., Berwick, J., 2015. Long-Latency Reductions in Gamma Power Predict Hemodynamic Changes That Underlie the Negative BOLD Signal. J. Neurosci. 35, 4641–4656. https://doi.org/10.1523/JNEUROSCI.2339-14.2015

Boorman, L., Kennerley, A.J., Johnston, D., Jones, M., Zheng, Y., Redgrave, P., Berwick, J., 2010. Negative Blood Oxygen Level Dependence in the Rat:A Model for Investigating the Role of Suppression in Neurovascular Coupling. J. Neurosci. 30, 4285–4294. https://doi.org/10.1523/JNEUROSCI.6063-09.2010

Cauli, B., Hamel, E., 2010. Revisiting the Role of Neurons in Neurovascular Coupling. Front Neuroenergetics 2. https://doi.org/10.3389/fnene.2010.00009

Cauli, B., Tong, X.-K., Rancillac, A., Serluca, N., Lambolez, B., Rossier, J., Hamel, E., 2004. Cortical GABA interneurons in neurovascular coupling: relays for subcortical vasoactive pathways. J Neurosci 24, 8940–8949. https://doi.org/10.1523/JNEUROSCI.3065-04.2004

Chang, J.C., Shook, L.L., Biag, J., Nguyen, E.N., Toga, A.W., Charles, A.C., Brennan, K.C., 2010. Biphasic direct current shift, haemoglobin desaturation and neurovascular uncoupling in cortical spreading depression. Brain 133, 996–1012. https://doi.org/10.1093/brain/awp338

Chow, B.W., Nuñez, V., Kaplan, L., Granger, A.J., Bistrong, K., Zucker, H.L., Kumar, P., Sabatini, B.L., Gu, C., 2020. Caveolae in CNS arterioles mediate neurovascular coupling. Nature 579, 106–110. https://doi.org/10.1038/s41586-020-2026-1

Dahlqvist, M.K., Thomsen, K.J., Postnov, D.D., Lauritzen, M.J., 2020. Modification of oxygen consumption and blood flow in mouse somatosensory cortex by cell-type-specific neuronal activity. J Cereb Blood Flow Metab 40, 2010–2025. https://doi.org/10.1177/0271678X19882787

Dirnagl, U., Lindauer, U., Villringer, A., 1993a. Nitric oxide synthase blockade enhances vasomotion in the cerebral microcirculation of anesthetized rats. Microvasc Res 45, 318–323. https://doi.org/10.1006/mvre.1993.1028

Dirnagl, U., Lindauer, U., Villringer, A., 1993b. Role of nitric oxide in the coupling of cerebral blood flow to neuronal activation in rats. Neurosci Lett 149, 43–46. https://doi.org/10.1016/0304-3940(93)90343-j

Dormanns, K., Brown, R.G., David, T., 2016. The role of nitric oxide in neurovascular coupling. J Theor Biol 394, 1–17. https://doi.org/10.1016/j.jtbi.2016.01.009

Drew, P.J., Shih, A.Y., Kleinfeld, D., 2011. Fluctuating and sensory-induced vasodynamics in rodent cortex extend arteriole capacity. Proceedings of the National Academy of Sciences 108, 8473–8478. https://doi.org/10.1073/pnas.1100428108

Dudek, F.E., 2020. Loss of GABAergic Interneurons in Seizure-Induced Epileptogenesis—Two Decades Later and in a More Complex World. Epilepsy Curr 20, 70S–72S. https://doi.org/10.1177/1535759720960464

Echagarruga, C.T., Gheres, K.W., Norwood, J.N., Drew, P.J., 2020. nNOS-expressing interneurons control basal and behaviorally evoked arterial dilation in somatosensory cortex of mice. Elife 9, e60533. https://doi.org/10.7554/eLife.60533

Eyre, B., Shaw, K., Sharp, P., Boorman, L., Lee, L., Shabir, O., Berwick, J., Howarth, C., 2022. The effects of locomotion on sensory-evoked haemodynamic responses in the cortex of awake mice. Sci Rep 12, 6236. https://doi.org/10.1038/s41598-022-10195-y

Faraci, F.M., Brian, J.E., 1994. Nitric oxide and the cerebral circulation. Stroke 25, 692–703. https://doi.org/10.1161/01.str.25.3.692

Fergus, A., Lee, K.S., 1997. GABAergic regulation of cerebral microvascular tone in the rat. J Cereb Blood Flow Metab 17, 992–1003. https://doi.org/10.1097/00004647-199709000-00009

Förstermann, U., Sessa, W.C., 2012. Nitric oxide synthases: regulation and function. Eur Heart J 33, 829–837, 837a–837d. https://doi.org/10.1093/eurheartj/ehr304

Gao, Y.-R., Ma, Y., Zhang, Q., Winder, A.T., Liang, Z., Antinori, L., Drew, P.J., Zhang, N., 2017. Time to wake up: Studying neurovascular coupling and brain-wide circuit function in the un-anesthetized animal. Neuroimage 153, 382–398. https://doi.org/10.1016/j.neuroimage.2016.11.069

Gebremedhin, D., Lange, A.R., Lowry, T.F., Taheri, M.R., Birks, E.K., Hudetz, A.G., Narayanan, J., Falck, J.R., Okamoto, H., Roman, R.J., Nithipatikom, K., Campbell, W.B., Harder, D.R., 2000. Production of 20-HETE and its role in autoregulation of cerebral blood flow. Circ Res 87, 60–65. https://doi.org/10.1161/01.res.87.1.60

Harder, D.R., Narayanan, J., Gebremedhin, D., 2011. Pressure-induced myogenic tone and role of 20-HETE in mediating autoregulation of cerebral blood flow. Am J Physiol Heart Circ Physiol 300, H1557–1565. https://doi.org/10.1152/ajpheart.01097.2010

Hariharan, A., Jing, Y., Collie, N.D., Zhang, H., Liu, P., 2019. Altered neurovascular coupling and brain arginine metabolism in endothelial nitric oxide synthase deficient mice. Nitric Oxide 87, 60–72. https://doi.org/10.1016/j.niox.2019.03.006

Harris, S., Bruyns-Haylett, M., Kennerley, A., Boorman, L., Overton, P.G., Ma, H., Zhao, M., Schwartz, T.H., Berwick, J., 2013. The Effects of Focal Epileptic Activity on Regional Sensory-Evoked Neurovascular Coupling and Postictal Modulation of Bilateral Sensory Processing. J Cereb Blood Flow Metab 33, 1595–1604. https://doi.org/10.1038/jcbfm.2013.115

Hays, C.C., Zlatar, Z.Z., Wierenga, C.E., 2016. The Utility of Cerebral Blood Flow as a Biomarker of Preclinical Alzheimer’s Disease. Cell Mol Neurobiol 36, 167– 179. https://doi.org/10.1007/s10571-015-0261-z

Hoiland, R.L., Caldwell, H.G., Howe, C.A., Nowak-Flück, D., Stacey, B.S., Bailey, D.M., Paton, J.F.R., Green, D.J., Sekhon, M.S., Macleod, D.B., Ainslie, P.N., 2020. Nitric oxide is fundamental to neurovascular coupling in humans. J Physiol 598, 4927–4939. https://doi.org/10.1113/JP280162

Hosford, P.S., Gourine, A.V., 2019. What is the key mediator of the neurovascular coupling response? Neurosci Biobehav Rev 96, 174–181. https://doi.org/10.1016/j.neubiorev.2018.11.011

Howarth, C., Mishra, A., Hall, C.N., 2021. More than just summed neuronal activity: how multiple cell types shape the BOLD response. Philos Trans R Soc Lond B Biol Sci 376, 20190630. https://doi.org/10.1098/rstb.2019.0630

Imig, J.D., Zou, A.P., Stec, D.E., Harder, D.R., Falck, J.R., Roman, R.J., 1996. Formation and actions of 20-hydroxyeicosatetraenoic acid in rat renal arterioles. Am J Physiol 270, R217–227. https://doi.org/10.1152/ajpregu.1996.270.1.R217

Iturria-Medina, Y., Sotero, R.C., Toussaint, P.J., Mateos-Pérez, J.M., Evans, A.C., Alzheimer’s Disease Neuroimaging Initiative, 2016. Early role of vascular dysregulation on late-onset Alzheimer’s disease based on multifactorial data-driven analysis. Nat Commun 7, 11934. https://doi.org/10.1038/ncomms11934

Kenny, A., Plank, M.J., David, T., 2018. The role of astrocytic calcium and TRPV4 channels in neurovascular coupling. J Comput Neurosci 44, 97–114. https://doi.org/10.1007/s10827-017-0671-7

Krawchuk, M.B., Ruff, C.F., Yang, X., Ross, S.E., Vazquez, A.L., 2020. Optogenetic assessment of VIP, PV, SOM and NOS inhibitory neuron activity and cerebral blood flow regulation in mouse somato-sensory cortex. J Cereb Blood Flow Metab 40, 1427–1440. https://doi.org/10.1177/0271678X19870105

Langford, D.J., Bailey, A.L., Chanda, M.L., Clarke, S.E., Drummond, T.E., Echols, S., Glick, S., Ingrao, J., Klassen-Ross, T., Lacroix-Fralish, M.L., Matsumiya, L., Sorge, R.E., Sotocinal, S.G., Tabaka, J.M., Wong, D., van den Maagdenberg, A.M.J.M., Ferrari, M.D., Craig, K.D., Mogil, J.S., 2010. Coding of facial expressions of pain in the laboratory mouse. Nat Methods 7, 447–449. https://doi.org/10.1038/nmeth.1455

Lecrux, C., Hamel, E., 2016. Neuronal networks and mediators of cortical neurovascular coupling responses in normal and altered brain states. Philos Trans R Soc Lond B Biol Sci 371. https://doi.org/10.1098/rstb.2015.0350

Lee, J., Stile, C.L., Bice, A.R., Rosenthal, Z.P., Yan, P., Snyder, A.Z., Lee, J.-M., Bauer, A.Q., 2021. Opposed hemodynamic responses following increased excitation and parvalbumin-based inhibition. J Cereb Blood Flow Metab 41, 841–856. https://doi.org/10.1177/0271678X20930831

Lee, L., Boorman, L., Glendenning, E., Christmas, C., Sharp, P., Redgrave, P., Shabir, O., Bracci, E., Berwick, J., Howarth, C., 2020. Key Aspects of Neurovascular Control Mediated by Specific Populations of Inhibitory Cortical Interneurons. Cereb Cortex 30, 2452–2464. https://doi.org/10.1093/cercor/bhz251

Leijenaar, J.F., van Maurik, I.S., Kuijer, J.P.A., van der Flier, W.M., Scheltens, P., Barkhof, F., Prins, N.D., 2017. Lower cerebral blood flow in subjects with Alzheimer’s dementia, mild cognitive impairment, and subjective cognitive decline using two-dimensional phase-contrast magnetic resonance imaging. Alzheimers Dement (Amst) 9, 76–83. https://doi.org/10.1016/j.dadm.2017.10.001

Lindauer, U., Megow, D., Matsuda, H., Dirnagl, U., 1999. Nitric oxide: a modulator, but not a mediator, of neurovascular coupling in rat somatosensory cortex. Am J Physiol 277, H799–811. https://doi.org/10.1152/ajpheart.1999.277.2.H799

Liu, X., Li, C., Falck, J.R., Roman, R.J., Harder, D.R., Koehler, R.C., 2008. Interaction of nitric oxide, 20-HETE, and EETs during functional hyperemia in whisker barrel cortex. Am J Physiol Heart Circ Physiol 295, H619–631. https://doi.org/10.1152/ajpheart.01211.2007

Lourenço, C.F., Laranjinha, J., 2021. Nitric Oxide Pathways in Neurovascular Coupling Under Normal and Stress Conditions in the Brain: Strategies to Rescue Aberrant Coupling and Improve Cerebral Blood Flow. Front Physiol 12, 729201. https://doi.org/10.3389/fphys.2021.729201

Ma, J., Ayata, C., Huang, P.L., Fishman, M.C., Moskowitz, M.A., 1996. Regional cerebral blood flow response to vibrissal stimulation in mice lacking type I NOS gene expression. Am J Physiol 270, H1085–1090. https://doi.org/10.1152/ajpheart.1996.270.3.H1085

Madisen, L., Mao, T., Koch, H., Zhuo, J., Berenyi, A., Fujisawa, S., Hsu, Y.-W.A., Garcia, A.J., Gu, X., Zanella, S., Kidney, J., Gu, H., Mao, Y., Hooks, B.M., Boyden, E.S., Buzsáki, G., Ramirez, J.M., Jones, A.R., Svoboda, K., Han, X., Turner, E.E., Zeng, H., 2012. A toolbox of Cre-dependent optogenetic transgenic mice for light-induced activation and silencing. Nat Neurosci 15, 793–802. https://doi.org/10.1038/nn.3078

Mayhew, J., Zheng, Y., Hou, Y., Vuksanovic, B., Berwick, J., Askew, S., Coffey, P., 1999. Spectroscopic analysis of changes in remitted illumination: the response to increased neural activity in brain. Neuroimage 10, 304–326. https://doi.org/10.1006/nimg.1999.0460

Mayhew, J.E., Askew, S., Zheng, Y., Porrill, J., Westby, G.W., Redgrave, P., Rector, D.M., Harper, R.M., 1996. Cerebral vasomotion: a 0.1-Hz oscillation in reflected light imaging of neural activity. Neuroimage 4, 183–193. https://doi.org/10.1006/nimg.1996.0069

Miettinen, R., Sirviö, J., Riekkinen, P., Laakso, M.P., Riekkinen, M., Riekkinen, P., 1993. Neocortical, hippocampal and septal parvalbumin- and somatostatin-containing neurons in young and aged rats: correlation with passive avoidance and water maze performance. Neuroscience 53, 367–378. https://doi.org/10.1016/0306-4522(93)90201-p

Mishra, A., Reynolds, J.P., Chen, Y., Gourine, A.V., Rusakov, D.A., Attwell, D., 2016. Astrocytes mediate neurovascular signaling to capillary pericytes but not to arterioles. Nat Neurosci 19, 1619–1627. https://doi.org/10.1038/nn.4428

Miyata, N., Taniguchi, K., Seki, T., Ishimoto, T., Sato-Watanabe, M., Yasuda, Y., Doi, M., Kametani, S., Tomishima, Y., Ueki, T., Sato, M., Kameo, K., 2001. HET0016, a potent and selective inhibitor of 20-HETE synthesizing enzyme. Br J Pharmacol 133, 325–329. https://doi.org/10.1038/sj.bjp.0704101

Mulligan, S.J., MacVicar, B.A., 2004. Calcium transients in astrocyte endfeet cause cerebrovascular constrictions. Nature 431, 195–199. https://doi.org/10.1038/nature02827

Oyekan, A.O., Youseff, T., Fulton, D., Quilley, J., McGiff, J.C., 1999. Renal cytochrome P450 omega-hydroxylase and epoxygenase activity are differentially modified by nitric oxide and sodium chloride. J Clin Invest 104, 1131–1137. https://doi.org/10.1172/JCI6786

Palop, J.J., Mucke, L., 2016. Network abnormalities and interneuron dysfunction in Alzheimer disease. Nat Rev Neurosci 17, 777–792. https://doi.org/10.1038/nrn.2016.141

Percie du Sert, N., Hurst, V., Ahluwalia, A., Alam, S., Avey, M.T., Baker, M., Browne, W.J., Clark, A., Cuthill, I.C., Dirnagl, U., Emerson, M., Garner, P., Holgate, S.T., Howells, D.W., Karp, N.A., Lazic, S.E., Lidster, K., MacCallum, C.J., Macleod, M., Pearl, E.J., Petersen, O.H., Rawle, F., Reynolds, P., Rooney, K., Sena, E.S., Silberberg, S.D., Steckler, T., Würbel, H., 2020. The ARRIVE guidelines 2.0: Updated guidelines for reporting animal research. PLoS Biol 18, e3000410. https://doi.org/10.1371/journal.pbio.3000410

Perrenoud, Q., Rossier, J., Férézou, I., Geoffroy, H., Gallopin, T., Vitalis, T., Rancillac, A., 2012. Activation of cortical 5-HT(3) receptor-expressing interneurons induces NO mediated vasodilatations and NPY mediated vasoconstrictions. Front Neural Circuits 6, 50. https://doi.org/10.3389/fncir.2012.00050

Poloyac, S.M., Zhang, Y., Bies, R.R., Kochanek, P.M., Graham, S.H., 2006. Protective effect of the 20-HETE inhibitor HET0016 on brain damage after temporary focal ischemia. J Cereb Blood Flow Metab 26, 1551–1561. https://doi.org/10.1038/sj.jcbfm.9600309

Salter, M., Duffy, C., Garthwaite, J., Strijbos, P.J., 1995. Substantial regional and hemispheric differences in brain nitric oxide synthase (NOS) inhibition following intracerebroventricular administration of N omega-nitro-L-arginine (L-NA) and its methyl ester (L-NAME). Neuropharmacology 34, 639–649. https://doi.org/10.1016/0028-3908(95)00036-6

Shabir, O., Pendry, B., Lee, L., Eyre, B., Sharp, P.S., Rebollar, M.A., Drew, D., Howarth, C., Heath, P.R., Wharton, S.B., Francis, S.E., Berwick, J., 2022. Assessment of neurovascular coupling and cortical spreading depression in mixed mouse models of atherosclerosis and Alzheimer’s disease. Elife 11, e68242. https://doi.org/10.7554/eLife.68242

Sharp, P.S., Ameen-Ali, K.E., Boorman, L., Harris, S., Wharton, S., Howarth, C., Shabir, O., Redgrave, P., Berwick, J., 2020. Neurovascular coupling preserved in a chronic mouse model of Alzheimer’s disease: Methodology is critical. J Cereb Blood Flow Metab 40, 2289–2303. https://doi.org/10.1177/0271678X19890830

Sharp, P.S., Shaw, K., Boorman, L., Harris, S., Kennerley, A.J., Azzouz, M., Berwick, J., 2015. Comparison of stimulus-evoked cerebral hemodynamics in the awake mouse and under a novel anesthetic regime. Scientific Reports 5, 12621. https://doi.org/10.1038/srep12621

Shaw, K., Boyd, K., Anderle, S., Hammond-Haley, M., Amin, D., Bonnar, O., Hall, C.N., 2021. Gradual Not Sudden Change: Multiple Sites of Functional Transition Across the Microvascular Bed. Front Aging Neurosci 13, 779823. https://doi.org/10.3389/fnagi.2021.779823

Singh, H., Cheng, J., Deng, H., Kemp, R., Ishizuka, T., Nasjletti, A., Schwartzman, M.L., 2007. Vascular cytochrome P450 4A expression and 20-hydroxyeicosatetraenoic acid synthesis contribute to endothelial dysfunction in androgen-induced hypertension. Hypertension 50, 123–129. https://doi.org/10.1161/HYPERTENSIONAHA.107.089599

Stefanovic, B., Schwindt, W., Hoehn, M., Silva, A.C., 2007. Functional uncoupling of hemodynamic from neuronal response by inhibition of neuronal nitric oxide synthase. J Cereb Blood Flow Metab 27, 741–754. https://doi.org/10.1038/sj.jcbfm.9600377

Sun, C.W., Alonso-Galicia, M., Taheri, M.R., Falck, J.R., Harder, D.R., Roman, R.J., 1998. Nitric oxide-20-hydroxyeicosatetraenoic acid interaction in the regulation of K+ channel activity and vascular tone in renal arterioles. Circ Res 83, 1069–1079. https://doi.org/10.1161/01.res.83.11.1069

Taniguchi, H., He, M., Wu, P., Kim, S., Paik, R., Sugino, K., Kvitsiani, D., Kvitsani, D., Fu, Y., Lu, J., Lin, Y., Miyoshi, G., Shima, Y., Fishell, G., Nelson, S.B., Huang, Z.J., 2011. A resource of Cre driver lines for genetic targeting of GABAergic neurons in cerebral cortex. Neuron 71, 995–1013. https://doi.org/10.1016/j.neuron.2011.07.026

Toth, P., Tarantini, S., Davila, A., Valcarcel-Ares, M.N., Tucsek, Z., Varamini, B., Ballabh, P., Sonntag, W.E., Baur, J.A., Csiszar, A., Ungvari, Z., 2015. Purinergic glio-endothelial coupling during neuronal activity: role of P2Y1 receptors and eNOS in functional hyperemia in the mouse somatosensory cortex. Am J Physiol Heart Circ Physiol 309, H1837–1845. https://doi.org/10.1152/ajpheart.00463.2015

Uhlirova, H., Kılıç, K., Tian, P., Thunemann, M., Desjardins, M., Saisan, P.A., Sakadžić, S., Ness, T.V., Mateo, C., Cheng, Q., Weldy, K.L., Razoux, F., Vandenberghe, M., Cremonesi, J.A., Ferri, C.G., Nizar, K., Sridhar, V.B., Steed, T.C., Abashin, M., Fainman, Y., Masliah, E., Djurovic, S., Andreassen, O.A., Silva, G.A., Boas, D.A., Kleinfeld, D., Buxton, R.B., Einevoll, G.T., Dale, A.M., Devor, A., 2016. Cell type specificity of neurovascular coupling in cerebral cortex. eLife Sciences 5, e14315. https://doi.org/10.7554/eLife.14315

Vazquez, A.L., Fukuda, M., Kim, S.-G., 2018. Inhibitory Neuron Activity Contributions to Hemodynamic Responses and Metabolic Load Examined Using an Inhibitory Optogenetic Mouse Model. Cereb Cortex 28, 4105–4119. https://doi.org/10.1093/cercor/bhy225

Verret, L., Mann, E.O., Hang, G.B., Barth, A.M.I., Cobos, I., Ho, K., Devidze, N., Masliah, E., Kreitzer, A.C., Mody, I., Mucke, L., Palop, J.J., 2012. Inhibitory interneuron deficit links altered network activity and cognitive dysfunction in Alzheimer model. Cell 149, 708–721. https://doi.org/10.1016/j.cell.2012.02.046

Vlasenko, O.V., Dovgan’, A.V., Maisky, V.A., Maznychenko, A.V., Pilyavskii, A.I., 2007. NADPH-Diaphorase reactivity and neurovascular coupling in the basal forebrain and motor cortex. Neurophysiology 39, 355–357. https://doi.org/10.1007/s11062-007-0056-z

Wang, Q., Kjaer, T., Jørgensen, M.B., Paulson, O.B., Lassen, N.A., Diemer, N.H., Lou, H.C., 1993. Nitric oxide does not act as a mediator coupling cerebral blood flow to neural activity following somatosensory stimuli in rats. Neurol Res 15, 33–36. https://doi.org/10.1080/01616412.1993.11740103

White, R.P., Hindley, C., Bloomfield, P.M., Cunningham, V.J., Vallance, P., Brooks, D.J., Markus, H.S., 1999. The effect of the nitric oxide synthase inhibitor L-NMMA on basal CBF and vasoneuronal coupling in man: a PET study. J Cereb Blood Flow Metab 19, 673–678. https://doi.org/10.1097/00004647-199906000-00011

Yuill, K.H., McNeish, A.J., Kansui, Y., Garland, C.J., Dora, K.A., 2010. Nitric oxide suppresses cerebral vasomotion by sGC-independent effects on ryanodine receptors and voltage-gated calcium channels. J Vasc Res 47, 93–107. https://doi.org/10.1159/000235964

